# Synergistic DNA and RNA binding of the Hox transcription factor Ultrabithorax coordinates splicing and shapes *in vivo* homeotic functions

**DOI:** 10.1101/2024.09.10.612310

**Authors:** Constanza Blanco, Wan Xiang, Panagiotis Boumpas, Maily Scorcelletti, Ashley Suraj Hermon, Jiemin Wong, Samir Merabet, Julie Carnesecchi

**Author notes:** The two first authors contribute equally and are listed in alphabetic order (Constanza Blanco, Wan Xiang).

## Abstract

The dual interaction of many transcription factors (TFs) with both DNA and RNA is an underexplored issue that could fundamentally reshape our understanding of gene regulation. We address this central issue by investigating the RNA binding activity of the *Drosophila* Hox TF Ultrabithorax (Ubx) in alternative splicing and morphogenesis. Relying on molecular and genetic interactions, we uncover a homodimerization-dependent mechanism by which Ubx regulates splicing. Notably, this mechanism enables the decoupling of Ubx-DNA and -RNA binding activity in splicing. We identify a critical residue for Ubx-RNA binding and demonstrate the essential role of Ubx-RNA binding ability for its homeotic functions. Overall, we uncover a unique mechanism for Ubx-mediated splicing and underscore the critical contribution of synergistic DNA/RNA binding for its morphogenetic functions. These findings advance our understanding of co-transcriptional regulation and highlight the significance of TF-DNA/RNA synergistic function in shaping gene regulatory networks in living organisms.

**Graphical abstract:** 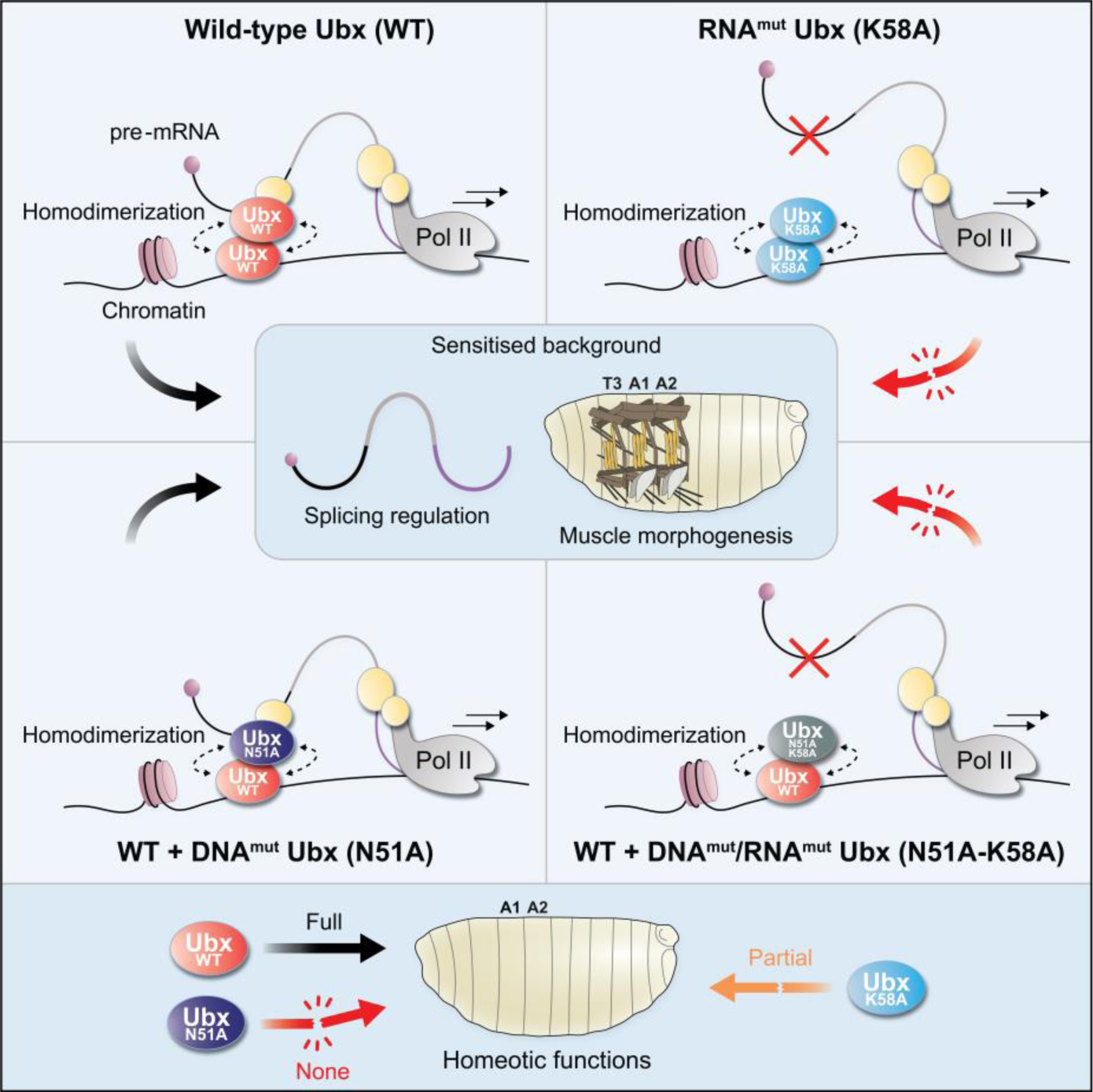

**Highlights:**

- Ubx homodimerization enables decoupling of DNA- and RNA-dependent splicing regulation
- The homeodomain K58 amino acid is critical for Ubx-RNA binding ability
- Ubx-RNA binding ability is essential for splicing regulation
- Dual DNA/RNA binding activities shape Ubx homeotic functions

## INTRODUCTION

Transcription factors (TFs) play a critical role in coordinating the development of multicellular organisms by regulating precise gene expression programs^1,2^. To this end, they recognise and bind to specific *cis*-regulatory DNA modules, thereby modulating the transcription of target genes in defined spatial and temporal contexts^3–5^. Beyond this conventional role in transcription, numerous studies have expanded our understanding of TF functions by revealing their key role in splicing^6–10^. Splicing is a critical process that generates transcript variability from a limited number of genes^11–13^. Alternative splicing, in particular, allows the production of multiple mRNA isoforms from a single pre-mRNA, thereby increasing transcriptome and proteome diversity^14–16^. Consequently, TF function in alternative splicing provides another regulatory layer of gene programs and highlights the TF multifaceted roles in gene expression^6,8^. Extensive research has revealed a tight functional and physical coupling between transcription and splicing^17–20^. Due to their comprehensive roles in both processes, TFs have emerged as key players in co-transcriptional alternative splicing^8,21,22^. However, unlike their well-characterised role in transcription, the mechanisms by which TFs influence splicing are elusive. Some TFs directly interact with the splicing machinery, while others indirectly influence splicing through the modulation of splicing factor expression or activity^6,10,23^. Additionally, certain TFs regulate splicing via their DNA binding activity and coordinate exon selection based on promoter identity^17,24^. All in all, how TFs coordinate both transcription and splicing to regulate precise morphogenetic networks in multicellular organisms remains largely unknown^8^.

Beyond their DNA binding ability, some TFs possess RNA binding abilities, adding another layer of complexity to their function^6,8,25–27^. For instance, Sox2-RNA binding contributes to cell pluripotency through splicing regulation^28^. Other TFs like SOX9 can bind RNA, yet its splicing activity is driven solely by DNA binding activity^23^. This variability in DNA and RNA binding activities underscores the remarkable complexity of TF functions. This complexity is exemplified by the Hox proteins, a family of homeodomain (HD)-containing TFs critical for anterior-posterior patterning during animal development and tissue homeostasis^29,30^. They are expressed along body polarity axes in cnidarians^31^ and bilaterians^29^ and orchestrate the development of various tissue types^29,32^. One member of the Hox family, the *Drosophila* TF Ultrabithorax (Ubx), has been extensively studied for its transcriptional function in morphogenesis, particularly in myogenesis and neurogenesis^5,33–35^. Notably, our recent work revealed that Ubx regulates gene expression at transcription and splicing levels in *Drosophila* cells and embryonic mesoderm^21^. Specifically, Ubx modulates co-transcriptional splicing through its DNA binding ability and interplay with active RNA Polymerase II (Pol II)^21^. Importantly, Ubx binds RNA *in vitro* and *in vivo*, yet the functional contribution of its RNA binding ability remains completely elusive. This raises the question of the significance of Ubx-DNA and -RNA binding abilities for its molecular and *in vivo* functions. Understanding this dual functionality of Ubx-DNA/RNA binding is thus crucial for unravelling the full molecular repertoire that shapes its gene regulatory programs during development.

Despite an important number of TFs possessing the dual capacity to bind DNA and RNA, it is still unclear if these binding capacities lead to two distinct functions, competition or synergy for shaping the gene regulatory networks. To address this issue, we investigated the dual DNA/RNA binding functionality of the Hox TF Ubx in alternative splicing. Our work demonstrated Ubx moonlighting functions beyond its DNA binding ability. We relied on the functional interplay between wild-type (WT) and mutant proteins to decouple the DNA- and RNA-dependent binding capability of Ubx and identified a new molecular mechanism by which Ubx regulates splicing through homodimerization. We generated and characterised the first Ubx-RNA binding mutant and demonstrated the critical function of Ubx-RNA binding in splicing regulation. Our work further unveiled the significance of Ubx-RNA binding activity for its homeotic function during *Drosophila* embryonic muscle development. In sum, our work reveals one of the mechanisms underpinning Ubx-splicing regulation via its synergistic DNA and RNA binding ability and dynamic homodimerization. Our results further emphasise the conservation of Hox-RNA binding ability across metazoans. Thus, integrating TF-DNA and TF-RNA binding activities is essential to elucidate the multiple TF functions orchestrating animal development.

## RESULTS

### Ubx exhibits *in vivo* moonlighting functions beyond its DNA-binding ability

Ubx binds RNA^21^ and yet, the functional implications of this association are unknown. To address this central question, we aimed to decouple and examine Ubx-DNA and -RNA binding activities on splicing and morphogenesis at the molecular and functional levels.

To this end, we focused on the *Drosophila* embryonic mesoderm, wherein Ubx modulates transcription and differential splicing^21^. Moreover, Ubx^WT^ physically and functionally interacts with splicing factors, notably with the spliceosome subunit snRNPU1-70K (U1-70K) for coordinating muscle development^35^. Similarly, Ubx^N51A^, a DNA binding mutant carrying a point mutation in the homeodomain (HD), physically interacts with splicing factors in the mesoderm, including U1-70K^35^. We thus hypothesised that this interaction is functional and tested it using the Ubx/U1-70K genetic interaction in embryonic muscle. A combined dose-reduction of Ubx and U1-70K disrupted proper muscle formation, as illustrated on a 3-dimensional (3D) lateral view of double heterozygous mutant embryos carrying one copy of the *Ubx*^1^ and *(snRNP)U1-70K^02107^* null mutant alleles (Figure 1A-C, stage 16). This disruption was particularly evident in the lateral transverse muscle number 3 (LT3) of abdominal A1 and A2 segments compared to control genotypes (100% penetrance, Figure 1A-C, Figure S1A-H). Having characterised the phenotype driven by the Ubx/U1-70K genetic interaction, we assessed the ability of Ubx transgenes to rescue the muscle alteration (Figure 1D-G). Using the GAL4/UAS system^36^, we expressed Ubx^WT^ or Ubx^N51A^ full-length proteins in the mesoderm (pan-mesodermal driver *mef2-GAL4*). We observed that both Ubx^WT^ and Ubx^N51A^ rescued the LT3 muscle alteration driven by the Ubx/U1-70K genetic interaction (Figure 1A, 1D-G). Of note, pan-mesodermal expression of Ubx^WT^ resulted in the typical homeotic transformation of thoracic T2-T3 muscles into abdominal ones, evidenced by the appearance of abdominal-specific ventral acute muscles (VA1, VA2, Figure 1D-E). Conversely, ectopic expression of the Ubx^N51A^ did not impact the identity of thoracic muscles, illustrating its inability to induce homeotic transformation by itself (Figure 1F-G). These data indicated that Ubx^N51A^ synergises with endogenous Ubx^WT^ activity and, by means, contributes to proper muscle development.

**Figure 1:**
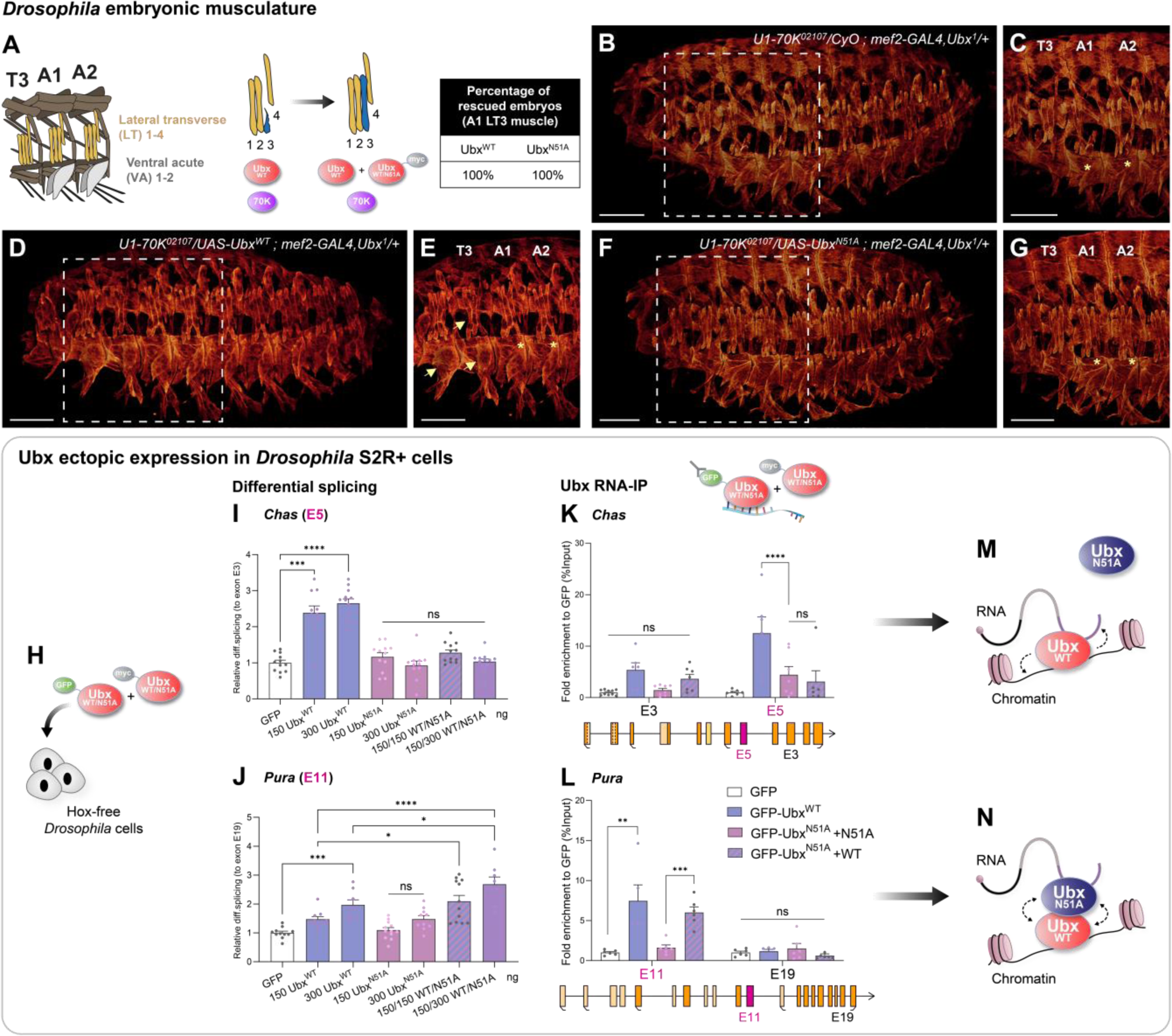
Ubx showcases moonlighting functions beyond DNA binding activity. (**A**) Schematic of (i) embryonic muscle patterns in thoracic T3 and abdominal A1, A2 segments (lateral transverse LT muscles yellow, ventral acute VA grey); (ii) muscle alteration driven by Ubx/U1-70K genetic interaction on LT3 muscle (blue) and rescue; (iii) rescue percentage of LT3 alteration upon pan-mesodermal Ubx^WT^ or Ubx^N51A^ (DNA^mut^) expression. (**B, D, F**) 3D projections of stage 16 embryos and (**C, E, G**) zooms (muscles stained with tropomyosin1). (**B-C**) Alteration of LT muscles in double heterozygous (*snRNP*)*U1-70K^02107^* with *Ubx^1^* embryos. Rescue experiments with pan-mesodermal expression (*mef2-GAL4*) of (**D-E**) Ubx^WT^ and (**F-G**) Ubx^N51A^. Yellow arrows indicate homeotic transformations of T2-T3 into an A1-like segment. Yellow stars indicate LT3. n=20 embryos per genotype. Scale bar 50 µm. (**H**-**J**) RTqPCR experiments showing the differential retention of exon cassettes over constitutive exons for (**I**) *Chas* and (**J**) *Pura* in *Drosophila* S2R+ cells expressing GFP control (white), Ubx^WT^ (blue), Ubx^N51A^ (pink) or both co-expressed (purple). Transfected plasmid quantity is indicated (ng). Ubx^N51A^ synergises with Ubx^WT^ splicing activity on *Pura*, regulated at the splicing level but not on *Chas* regulated at transcription and splicing levels. (**K-L**) Nuclear RNA-immunoprecipitation (RIP-RTqPCR) in cells expressing GFP, GFP-Ubx^WT^ and GFP-Ubx^N51A^ with myc-Ubx^WT^ or myc-Ubx^N51A^ on constitutive and differentially spliced exons of (**K**) *Chas* and (**L**) *Pura*. Values are relative enrichment over GFP relative to input. Ubx^N51A^ associates with the cassette exon of *Pura* when co-expressed with Ubx^WT^. Schematic of *Chas* and *Pura* gene architecture highlighting exons (cassette exons in pink) with numbers (E) following JunctionSeq annotation^21^. Black arrows represent transcription directionality. Alternative transcription start (TSS) and termination (TTS) sites are in brackets. Bars represent mean ±SEM for 3-4 biological replicates. Statistics by one-way ANOVA (\**P*< 0.05, \*\**P*< 0.01, \*\*\**P*< 0.001, ****P< 0.0001, ns = non-significant). (**M-N**) Schematic of synergistic splicing activity of Ubx^WT^/Ubx^N51A^ via dimerization on *Pura* but not on *Chas.* **See also Figures S1-S2**.

Next, we characterised the minimal domain required for rescuing the Ubx/U1-70K genetic interaction. Several truncated Ubx derivatives were employed for rescue experiments, showing that the Ubx HD contributes significantly (80%) to rescuing muscle alteration driven by the Ubx/U1-70K genetic interaction (Figure S1I-O). Additional *in vitro* experiments confirmed the physical interaction between Ubx HD and U1-70K proteins (Figure S1P-R).

In conclusion, these results highlight a functional interplay between Ubx^WT^ and Ubx^N51A^ in regulating muscle formation via U1-70K interaction. Importantly, these data indicate that Ubx functions in myogenesis extend beyond its DNA binding ability.

### Ubx-splicing activity is not solely determined by its DNA binding capacity

Our *in vivo* data suggest that Ubx molecular functions do not uniquely rely on its DNA binding ability. Although Ubx^N51A^ cannot coordinate transcription, it could regulate splicing when co-expressed with Ubx^WT^, thereby rescuing the Ubx/U1-70K dose reduction *in vivo*. To test this idea, we evaluated the splicing activity of Ubx^N51A^ in a Hox-free *Drosophila* cell system, wherein WT and mutant Ubx co-expression and levels can be tightly controlled (Figure 1H).

We analysed the Ubx splicing activity on target genes identified previously^21^, including *Chascon* (*Chas*), regulated at both transcriptional and splicing levels, three genes regulated solely at the splicing level, *Puratrophin* (*Pura*), *polyA binding protein* (*pAbp*), *Dunce* (*Dnc*), and *Rad, Gem/Kir family member 1* (*Rgk1*) exhibiting an intermediary profile with Ubx^WT^ influencing differential splicing, but also transcription only at high Ubx expression level (Figure 1I-J, Figure S2A-N). Ubx^N51A^ failed to induce differential splicing and expression of *Chas,* whether expressed alone or with Ubx^WT^ (Figure 1I, Figure S2A-B). Conversely, Ubx^N51A^ co-expressed with Ubx^WT^ promoted the retention of cassette exons of Ubx-target genes regulated exclusively at the splicing level (while their expression remained unchanged). This was the case for *pAbp*, *Pura*, *Rgk1* and *Dnc* (Figure 1J, Figure S2H, S2K, S2N). Importantly, this effect was dose-dependent, as increased Ubx^N51A^ level (with constant Ubx^WT^ level) led to higher exon inclusion for *pAbp*, *Pura* and *Dnc* (Figure 1J, Figure S2H, S2N). The effect was moderate for *Rgk1*, for which Ubx^WT^ regulates its expression level at higher doses (Figure S2I-K). These results strongly suggest that Ubx employs distinct molecular mechanisms to regulate exclusively spliced genes (*Pura*) compared to the gene regulated at both transcription and splicing levels (*Chas*). Noteworthy, these mechanisms depend on Ubx protein concentration.

Both Ubx^WT^ and Ubx^N51A^ bind RNA *in vitro*^21^. However, *in vivo*, only Ubx^WT^, and not Ubx^N51A^, associates with differentially spliced exons of target transcripts (Figure 1K-L, Figure S2O-Q). We thus assessed the capacity of Ubx^N51A^ to bind RNA when co-expressed with Ubx^WT^ *in vivo*. Strikingly, nuclear Ubx-RNA immunoprecipitation in *Drosophila* S2R+ cells showed that Ubx^N51A^ is enriched on RNA in the presence of Ubx^WT^. Specifically, Ubx^N51A^-RNA association was strongly detected on exons differentially spliced upon Ubx^WT^/Ubx^N51A^ co-expression (*Pura*, *pAbp)*, with moderate association on *Rgk1* and absence on *Chas* (Figure 1K-L, Figure S2P-Q). These results are consistent with the synergistic splicing activity of Ubx^WT^/Ubx^N51A^ observed on differentially spliced transcripts (*Pura*) but not on Ubx-target genes regulated at both transcription and splicing levels (*Chas*) (Figure 1I-J). While both gene categories imply Ubx-RNA binding events coordinating splicing, two distinct mechanisms seem at play: the target genes regulated at both transcription and splicing levels largely depend on Ubx-DNA binding ability to regulate splicing, which cannot be dissociated from its RNA binding activity (Figure 1M). This entanglement challenges the decoupling of Ubx-DNA and -RNA binding activities. Contrarywise, the target genes regulated exclusively at the splicing level rely on distinct DNA-dependent and RNA-dependent Ubx molecular functions. For this gene class, we propose that Ubx^WT^ and Ubx^N51A^ dimerize, thereby facilitating the Ubx^N51A^-RNA association and splicing regulation (Figure 1N).

In sum, our data imply that Ubx splicing activity broadly depends on RNA binding and, thus, underpins moonlighting functions for Ubx beyond its DNA binding activity. Our results further suggest that Ubx-target genes exclusively spliced rely on an additional Ubx dimerization mechanism to promote splicing. As 70% of spliced Ubx-targets are exclusively differentially spliced in the embryonic mesoderm^21^, this mechanism could be crucial for Ubx morphogenetic function during muscle development.

### Ubx nuclear distribution facilitates its DNA-independent splicing function

We decoupled Ubx-DNA and Ubx-RNA binding requirements on exclusively spliced genes, thanks to the Ubx^WT/^Ubx^N51A^ coordinated action. As most differentially spliced Ubx-targets in embryonic mesoderm are exclusively spliced^21^, we extended our investigation to elucidate the mechanism regulating this important gene class, relying on the Ubx^WT^/Ubx^N51A^ interaction. Based on the Hox dimerization ability^37,38^, we hypothesised that the Ubx^WT^/Ubx^N51A^ physical interaction relocates Ubx^N51A^ close to its target transcripts, enabling Ubx^N51^-RNA binding and splicing regulation in a DNA binding-independent fashion.

To test this hypothesis, we analysed the Ubx dimerization potential *in vitro,* showing that Ubx^WT^ and Ubx^N51A^ interact and that the HD is sufficient for Ubx dimerization (Figure S3A-B). Native protein-DNA and protein-RNA interaction assays confirmed the Ubx dimerization potential on nucleic acids (EMSA, Figure S3C). Notably, Ubx-RNA binding does not rely on the nature of the RNA *in vitro*^21^. Thus, we used *Chas* probes due to their suitability for both denaturing and native assays^21^. To detect multimeric protein/DNA/RNA complexes, we performed UV-crosslinking assays with DNA, RNA and purified MBP-Ubx^WT^ (80kDa) and his-Ubx^N51A^ (45kDa) distinguished by their size under denaturing conditions (Figure S3D). Noteworthy, overlapping RNA and DNA signals over the Ubx^WT^ monomer suggest that one Ubx molecule can contact RNA and DNA simultaneously. Although we cannot undoubtedly distinguish the DNA/RNA overlap for Ubx dimer due to fuzzy DNA signal, we distinctly observed homo- and heterodimers Ubx^WT^/Ubx^N51A^ on RNA. Altogether, these results strongly support the likelihood of monomeric and dimeric protein/DNA/RNA complexes.

Next, we tested the interaction in *Drosophila* S2R+ cells by co-immunoprecipitation under native conditions (Figure 2A). Strikingly, we observed a strong interaction between Ubx^WT^ proteins, while the Ubx^WT^/Ubx^N51A^ association was strongly reduced (Figure 2A). In contrast, the Ubx^WT^/Ubx^N51A^ interaction was stronger upon formaldehyde crosslinking (Figure 2A), which allows the capture of transient protein-protein interactions^39^. Thus, the interaction prevalence could sufficiently confine the Ubx protein close to its target transcripts. We tested this idea using live imaging with Fluorescence Recovery After Photobleaching (FRAP) experiments, aiming to analyse the dynamic behaviour of GFP-Ubx^N51A^ (DNA^mut^) proteins in the presence of Ubx^WT^ (Figure 2B-E). The nuclear dynamics uncovered by FRAP are distinct for Ubx^N51A^ and Ubx^WT^ proteins^21^. Ubx^N51A^ is mainly present in diffusible fractions and is associated with a fast recovery rate (fast molecules) due to its inability to bind nucleic acids *in vivo*. In contrast, Ubx^WT^ is present in immobile or slower fractions, and its recovery rate is slower due to its nucleic acid binding ability (Figure 2B-E). When co-expressed with Ubx^WT^, Ubx^N51A^ protein localisation and dynamics were significantly changed. This was illustrated by a decrease in the fast mobile fraction (less diffusive molecules) and half-time recovery of the bleached area (i.e., slower molecules) (Figure 2C-E). This strongly suggested that Ubx^N51A^ relocalisation facilitates RNA association, thereby slowing down the Ubx^N51A^ recovery rate. Notably, we did not observe Ubx^N51A^ enrichment on the chromatin upon co-expression with Ubx^WT,^ emphasising the dynamic nature of the Ubx^WT^/Ubx^N51A^ interaction (Figure S3E-J, Ubx bound regions^21^).

**Figure 2:**
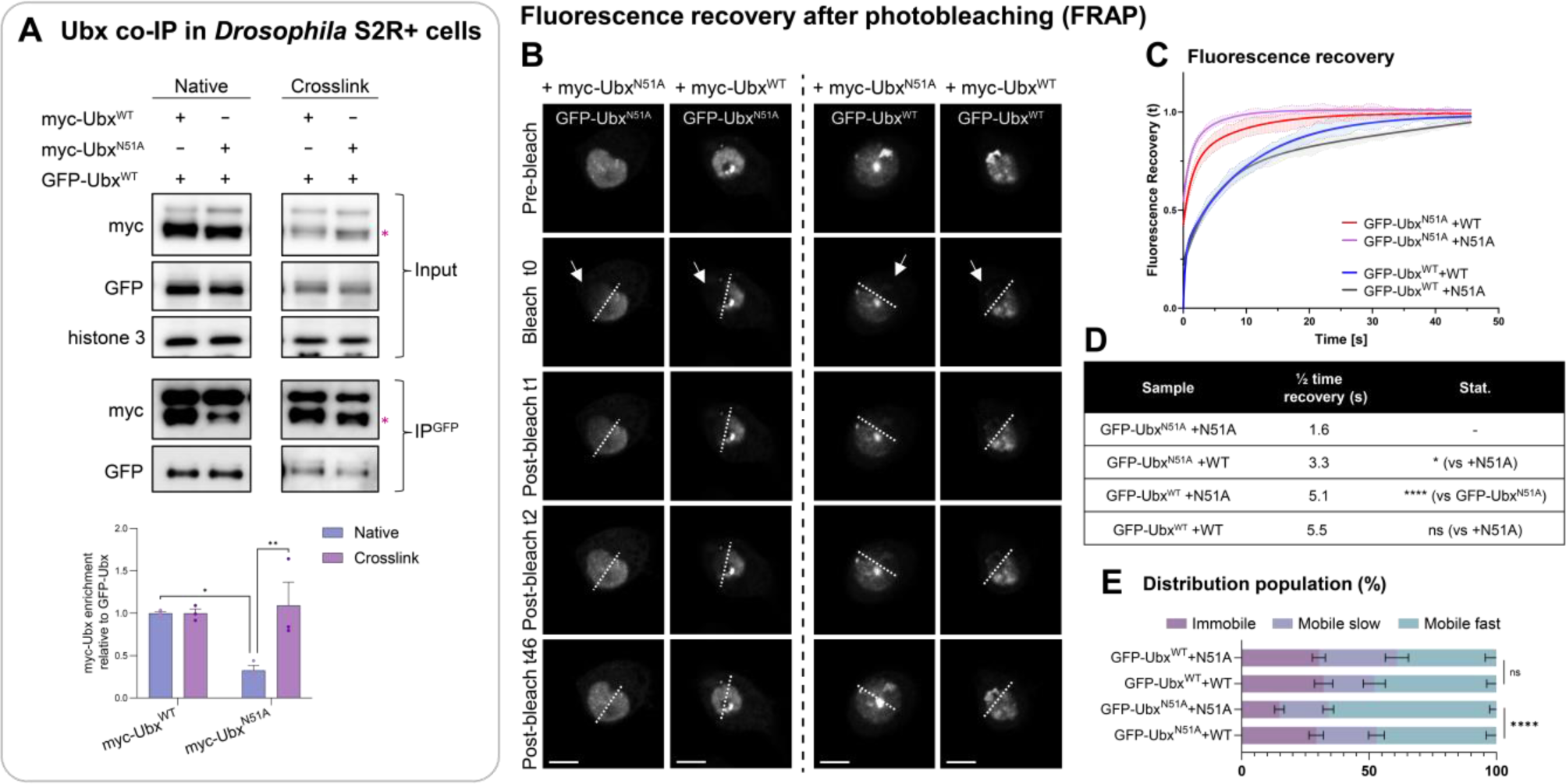
Ubx-DNA binding ability and dynamic homodimerization shape its nuclear dynamics. (**A**) Co-immunoprecipitation of myc-Ubx (pink star) derivatives with GFP-Ubx^WT^ with or without formaldehyde crosslink. Western blots were probed with indicated antibodies. The input is shown with histone 3 as a loading control. Quantification of relative enrichments to GFP-Ubx^WT^ pull-down showed a robust Ubx^N51A^ interaction upon crosslink compared to native conditions. n=3 biological replicates. Bars represent mean ±SEM. Statistics by one-way ANOVA (*P< 0.05, **P< 0.01). (**B**) Representative pictures of cells expressing GFP-Ubx^WT/N51A^ in the presence of myc-Ubx^WT/N51A^ during FRAP assay (pre-bleached, bleached (t0) and post-bleached images t1, t2, t46). White arrows indicate bleached areas. Scale bar 5 µm. (**C**) Normalised curves of fluorescence recovery (t) after photobleaching related to time (s, second) and (**D**) calculated half-time recoveries. Modelling follows a bi-exponential model^21^. (**E**) Distribution of Ubx populations: immobile represents the fraction stably loaded onto chromatin; slow mobile, intermediate interactions or scanning behaviour and fast mobile, the transient interaction and diffusive Ubx molecules. Means ±SEM are shown. One-way ANOVA and Chi^2^-test were applied respectively for half-time recoveries and population distribution. (*P< 0.05, ****P< 0.0001, ns = non-significant). n=25-32 nuclei per sample. **See also Figure S3**.

Altogether, these data revealed that the transient interaction between Ubx^WT^ and Ubx^N51A^ is sufficient to reshape Ubx^N51A^ nuclear dynamics. We propose that the interaction prevalence confines Ubx proteins close to Ubx-target transcripts, thus facilitating RNA binding and DNA-independent splicing regulation.

### The K58 residue of the HD is essential for Ubx-RNA but not Ubx-DNA binding ability

Overall, our results strongly suggest that Ubx-RNA binding is essential for regulating differential splicing globally. To unravel this mechanism, we investigated the Ubx-RNA binding interface to identify amino acids critical for Ubx-RNA association.

We characterised Ubx-RNA binding by *in vitro* UV-crosslinking assays on distinct Ubx-RNA targets (*Chas, Pura, pAbp, Rgk1*) bound by Ubx^WT^ and Ubx^N51A^. Using various Ubx^WT^ protein derivatives, we found that HD is the minimal domain required for Ubx to bind RNA, irrespective of the RNA nature. In line with the HD ability to rescue Ubx/U1-70K genetic interaction *in vivo*, the N-terminal and C-terminal counterparts did not bind RNA without the HD (Figure 3A-F, Figure S4A-F). Notably, the HD combined with a short C-terminal motif termed UBDA is sufficient to confer the complete Ubx-RNA binding potential *in vitro* (Figure S3G-I, Figure S4G-I). Additional dissection of the HD revealed that the third α-helix is the main anchor for Ubx-RNA interaction (Figure 3J-L, Figure S4J-L).

**Figure 3:**
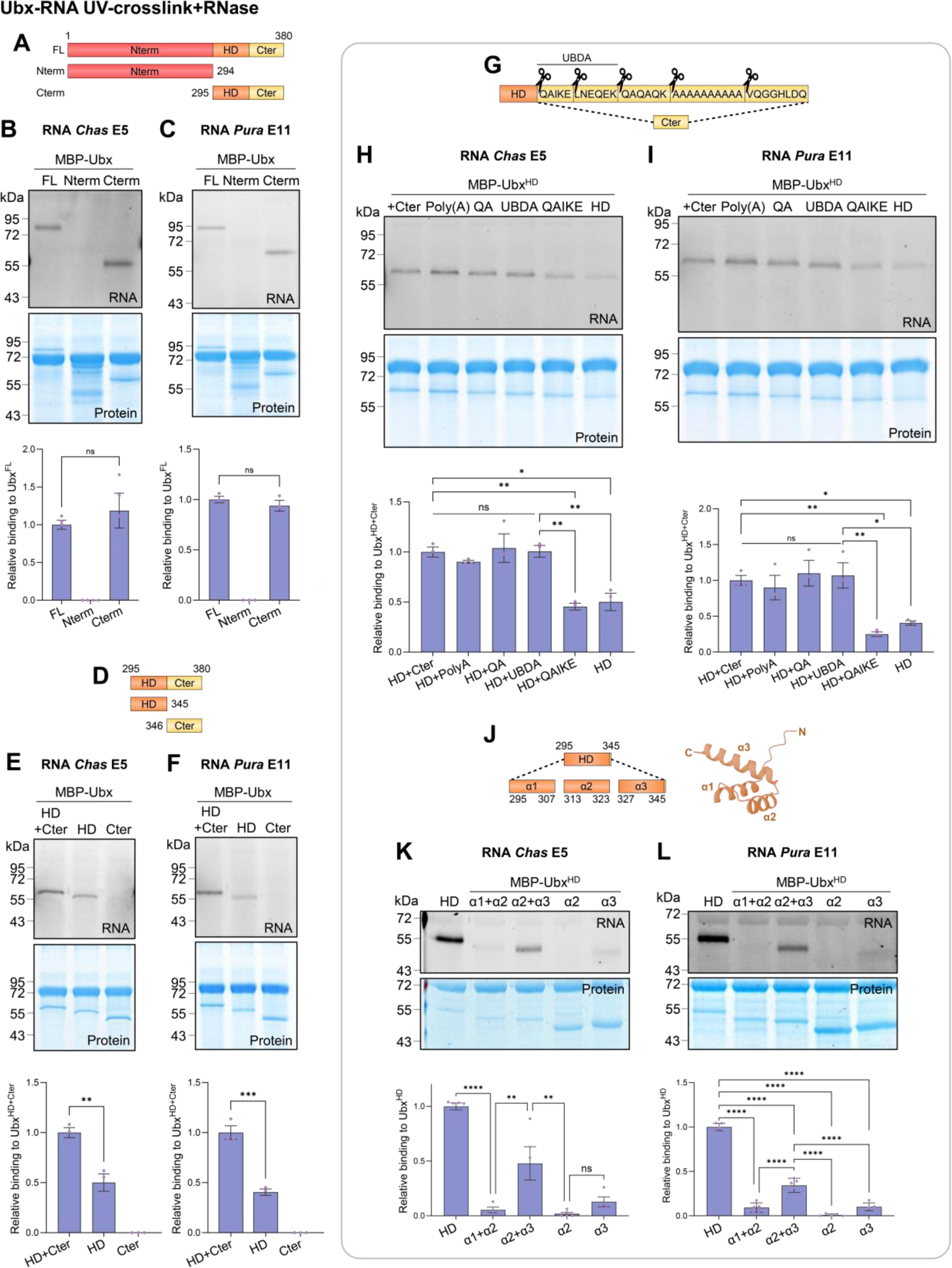
The HD α3-helix is central for Ubx-RNA binding, while the UBDA motif confers affinity. (**A, D, G, J**) Cartoon of domains and motifs dissected by *in vitro* UV-crosslinking assays. (**B-C, E-F, H-I, K-L**) Protein-RNA interaction followed by UV-crosslink and RNase digestion performed *in vitro* with purified proteins his-MBP-Ubx derivatives as indicated on (**B, E, H, K**) *Chas* Exon E5 or (**C, F, I, L**) *Pura* Exon E11 RNA probes. Cy3-UTP (RNA) signal detected interactions, Coomassie reveals the protein content (BSA is detected at 70kDa). Molecular marker is indicated. (**J**) HD structure generated with *AlphaFold*^50^. Quantification of relative RNA binding of Ubx derivatives compared to (**B-C**) full-length (FL), (**E-F, H-I**) HD+Cter or (**K-L**) HD for each RNA probe normalised to Coomassie (Protein). n=3 biological replicates. Bars represent mean ±SEM. Statistical test by one-way ANOVA (\**P*< 0.05, \*\**P*< 0.01, \*\*\**P*< 0.001, ****P< 0.0001, ns = non-significant). **See also Figure S4**.

As the α3-helix sequence is evolutionarily conserved for DNA binding^40^, we generated several HD mutants of conserved amino acids (Figure 4A, Figure S5A-B)^41^. Strikingly, when amino acids in the α3-helix core were mutated, HD-RNA and -DNA binding was strongly weakened, or protein folding was severely impaired (F49Q and R53A, Figure S5A-B). Thus, we targeted amino acids downstream of the DNA binding core in the α3-helix C-terminal extremity (Figure 4A, Figure S5C-K). We focused on two positively charged amino acids, the lysine K57 and K58, for which Ubx retained robust DNA binding ability when mutated into alanine (K57A, K58A, Figure S5C-D). Remarkably, the K57A mutation appeared to reduce Ubx binding on DNA (Figure S5D). Next, we evaluated the *in vitro* RNA binding ability of these Ubx mutants. While mutating lysine K57 did not impact Ubx-RNA binding capability, mutation of lysine K58 strongly and significantly reduced Ubx-RNA binding ability (Figure 4B-C, Figure S5E, S5H-I). This was observed for full-length and HD derivatives on different RNA probes and confirmed by electrophoretic mobility shift assay (Figure S5E, S6A-B). Having identified the lysine K58 as a critical residue for Ubx-RNA binding, various mutations (K58L, K58E, K58R, K58A) were tested, and alanine (K58A) was identified as the critical amino acid substitution impairing Ubx-RNA binding while retaining DNA binding ability (Figure S6A-E).

**Figure 4:**
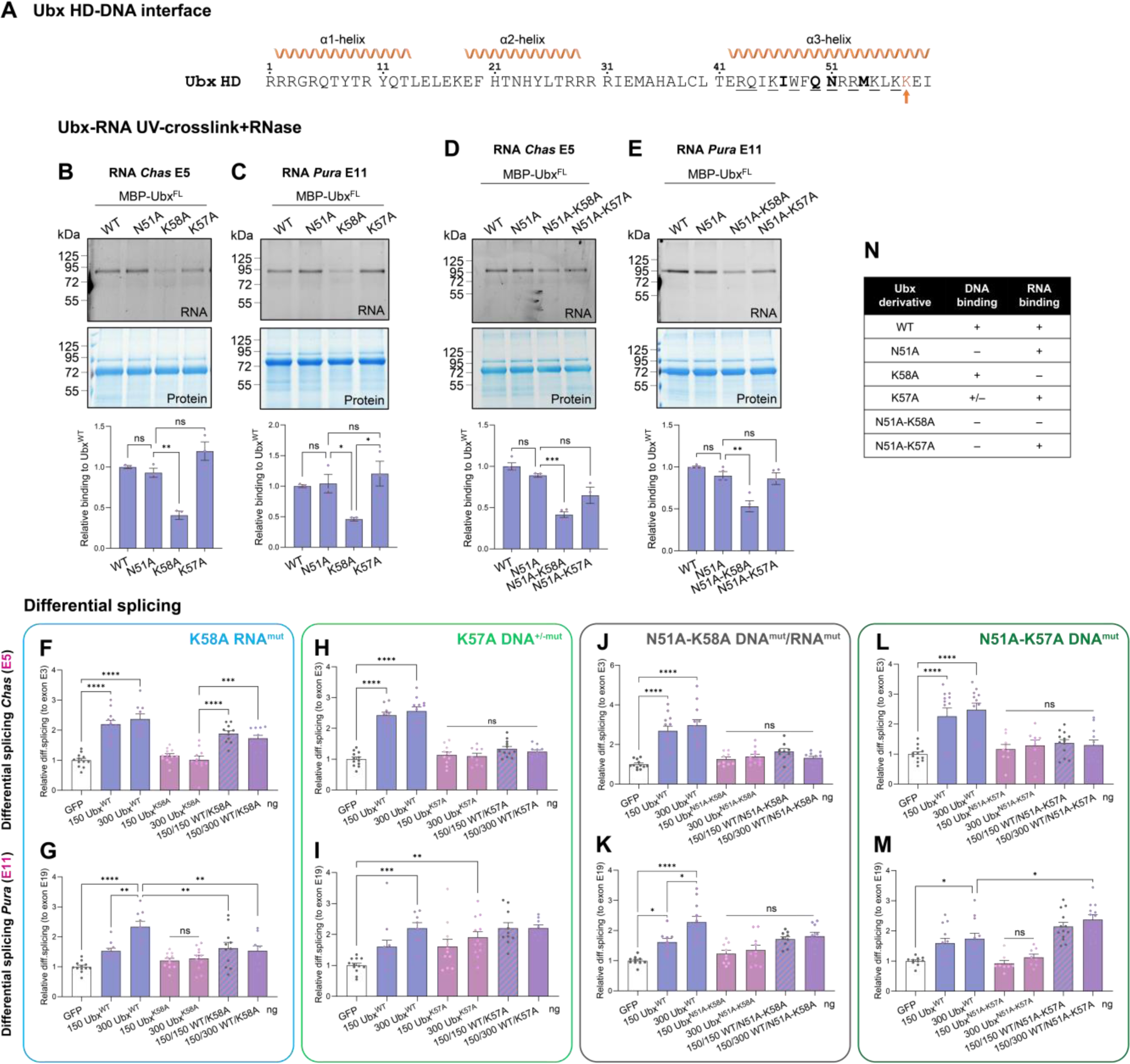
The K58 amino acid is essential for Ubx-RNA binding ability and splicing activity. (**A**) Ubx HD sequence and helices. Amino acids: In bold, interacting with the DNA major groove; underlined, contacting the DNA phosphate moiety; K58 (orange arrow). (**B-E**) UV-crosslinking assays with purified proteins his-MBP-Ubx as indicated on (**B, D**) *Chas* or (**C, E**) *Pura* RNA probes for (**B-C**) single or (**D-E**) combinatorial Ubx mutants. Cy3-UTP (RNA) signal detected interactions, Coomassie reveals the protein content (BSA at 70kDa). Molecular marker is indicated. The mutation K58A impacts Ubx-RNA binding ability *in vitro* but not the K57A mutation, alone or combined with N51A (DNA^mut^). n=3 biological replicates. Bars represent mean ±SEM of ratio Cy3-RNA over Coomassie-Protein. (**F-M**) Differential retention of exon cassettes over constitutive exons for *Chas* (**F, H, J, L**) and *Pura* (**G, I, K, M**) in *Drosophila* S2R+ cells expressing GFP control (white), Ubx^WT^ (blue), (**F-G**) Ubx^K58A^ (RNA^mut^, blue), (**H-I**) Ubx^K57A^ (DNA^+/-mut^, green) or (**J-K**) Ubx^N51A-K58A^ (DNA^mut^/RNA^mut^, black) or (**L-M**) Ubx^N51A-K57A^ (DNA^mut^, dark green) (pink) or both co-expressed (purple). Transfected plasmid quantity is indicated (ng). While Ubx^N51A-K57A^ synergises with Ubx splicing activity on *Pura* mRNA (only regulated at the splicing level), Ubx^K58A^ and Ubx^N51A-K58A^ mutants fail to promote Ubx splicing activity, demonstrating the importance of Ubx-RNA binding in its splicing activity. n=4 biological triplicate. Bars are mean ±SEM. Statistics by one-way ANOVA (\**P*< 0.05, \*\**P*< 0.01, \*\*\**P*< 0.001, \*\*\*\**P*< 0.0001, ns = non-significant). (**N**) Summary of the mutant transgenes. **See also Figures S5-S12**.

Subsequently, we evaluated the RNA and DNA binding ability of Ubx mutants *in vivo*. To this end, we performed nuclear Ubx-RNA immunoprecipitation on Ubx-target transcripts (*Chas, Pura, pAbp, Rgk1*) upon expression of Ubx derivatives in the Hox-free *Drosophila* S2R+ cells. Ubx^K58A^ exhibited a significantly reduced association with RNA *in vivo* compared to the Ubx^WT^ protein (Figure S7A-E). This confirmed that the K58A mutation impairs Ubx-RNA association on Ubx-target differentially spliced RNA exons. We further assessed the chromatin occupancy of Ubx mutants on Ubx-bound genomic regions (gene body for *Chas, Pura, pAbp, Rgk1* and intergenic enhancers *teashirt* (*tsh)* and *decapentaplegic* (*dpp*), Figure S7F-K). Except for the *tsh* enhancer (1.5 fold decrease), the Ubx^K58A^ chromatin occupancy was analogous to Ubx^WT^ (Figure S7F-K). In contrast, the K57A mutation did not circumvent the Ubx-RNA association but partly impacted Ubx-DNA binding ability *in vivo*, acting like a moderate DNA binding mutant (Figures S7L-V).

Altogether, these results showed that the K58A mutation specifically impairs Ubx-RNA but not -DNA binding *in vitro* and *in vivo*.

### The Ubx-RNA binding ability is essential for splicing activity

Relying on our identification of a Ubx-RNA binding mutant retaining DNA binding capacity, we sought to evaluate the Ubx-RNA binding requirement in splicing regulation.

We analysed differentially spliced Ubx-target transcripts upon Ubx^K58A^ expression in *Drosophila* S2R+ cells (Figure 4F-G, Figure S8A-N). As expected, the K58A mutation broadly circumvents Ubx splicing activity both on Ubx-target genes regulated at transcription and splicing levels (*Chas, Rgk1*) or exclusively spliced (*Pura, pAbp, Dnc*). Moreover, unlike the N51A mutation, Ubx^K58A^ (RNA^mut^) cannot synergise with Ubx^WT^ splicing activity when co-expressed (Figure 4G, Figure S8H, S8N). As a control, Ubx^K57A^ (DNA^+/-mut^) cannot promote transcription but promotes moderate splicing and synergises with Ubx^WT^ splicing activity on gene solely spliced, similar to Ubx^N51A^ (DNA^mut^) (*Pura, pAbp, Dnc*, Figure 4H-I, Figure S9A-N). These data reinforced that the K58A mutation specifically disrupts Ubx-RNA binding ability and, consequently, Ubx splicing activity.

Furthermore, we utilised the Ubx^WT^/Ubx^N51A^ synergistic activity observed on the gene class exclusively differentially spliced (*Pura, pAbp, Dnc*) to decouple Ubx-DNA from Ubx-RNA binding activity. We generated a mutant carrying both N51A and K58A mutations, thereby abolishing its whole nucleic acid binding ability (Figure 4D-E, Figure S5F-G, S5J-K). We analysed the Ubx splicing activity upon Ubx^WT^ co-expression. In sharp contrast to Ubx^N51A^ (DNA^mut^), the combinatorial mutant Ubx^N51A-K58A^ (DNA^mut^/RNA^mut^) failed to synergise with the Ubx^WT^ splicing activity (Figure 4J-K, Figure S10A-N). Additional Ubx-RNA immunoprecipitation confirmed that the Ubx^N51A-K58A^ mutant fails to associate with RNA exons in the presence of Ubx^WT^ (Figure S7A-E). In contrast, Ubx^N51A-K57A^ (DNA^mut^) retained the capacity to bind RNA when co-expressed with Ubx^WT^ and to promote exon inclusion in Ubx-target transcripts solely spliced (Figure 4L-M, Figure S7L-P, S11A-N). Importantly, this is not due to loss of dimerization potential as both Ubx^N51A-K58A^ and Ubx^N51A-K57A^ interact with Ubx^WT^ *in vitro* and *in vivo* (Figure S12A-D). Precisely, the Ubx-RNA association reflects the ability of Ubx^N51A^ and Ubx^N51A-K57A^ to synergise with Ubx^WT^ splicing activity on exclusively differentially spliced targets. In sharp contrast, Ubx^N51A-K58A^ loses both RNA binding and splicing activity when co-expressed with Ubx^WT^ protein, indicating that the K58A mutation specifically impairs Ubx-RNA binding and splicing activity. Notably, genes regulated at both transcription and splicing levels (*Chas, Rgk1* at high Ubx level) are equally impacted by the loss of DNA and RNA binding ability.

In sum, our results demonstrated that the Ubx-RNA binding is essential for Ubx splicing activity. Importantly, Ubx-DNA and Ubx-RNA binding requirements can be functionally decoupled on Ubx-target genes uniquely differentially spliced.

### Ubx-RNA binding activity contributes to muscle formation

Having demonstrated that Ubx-RNA binding ability regulates splicing, the functional relevance of the Ubx-RNA association during development was questioned.

We generated *Drosophila* transgenes expressing RNA- (Ubx^K58A^), DNA- (Ubx^K57A^, Ubx^N51A-K57A^) or combinatorial (Ubx^N51A-K58A^) binding mutants complementing our transgenic toolkit (Ubx^WT^, Ubx^N51A^) (Figure 4N, 5A-H). We utilised myogenesis as a biological context, with the sensitised Ubx/U1-70K genetic background (*Ubx^1^* and *U1-70K^02107^* double heterozygous mutant embryos) amenable to reveal muscle morphogenetics defects independent of Ubx-DNA binding ability (Figure 1A-G, LT3 alteration). As observed earlier, mesoderm-specific expression of Ubx-DNA binding mutants (Ubx^K57A^ or Ubx^N51A-K57A^) in Ubx/U1-70K genetic interaction context rescued the A1-A2 LT3 muscle alteration in embryos (Figure 5C-D, 5G-H). Conversely, expression of RNA binding mutants (Ubx^K58A^ or Ubx^N51A-K58A^) failed to rescue the A1-A2 LT3 muscle alteration (Figure 5A-B, 5E-F). These observations were supported by quantification of the A1-A2 LT3 muscle thickness, which showed significant changes compared to the control genotypes for RNA-binding mutant transgenes and muscle thickness rescue for DNA-binding mutant proteins (Figure 5I-J, Figure S13A-I).

**Figure 5:**
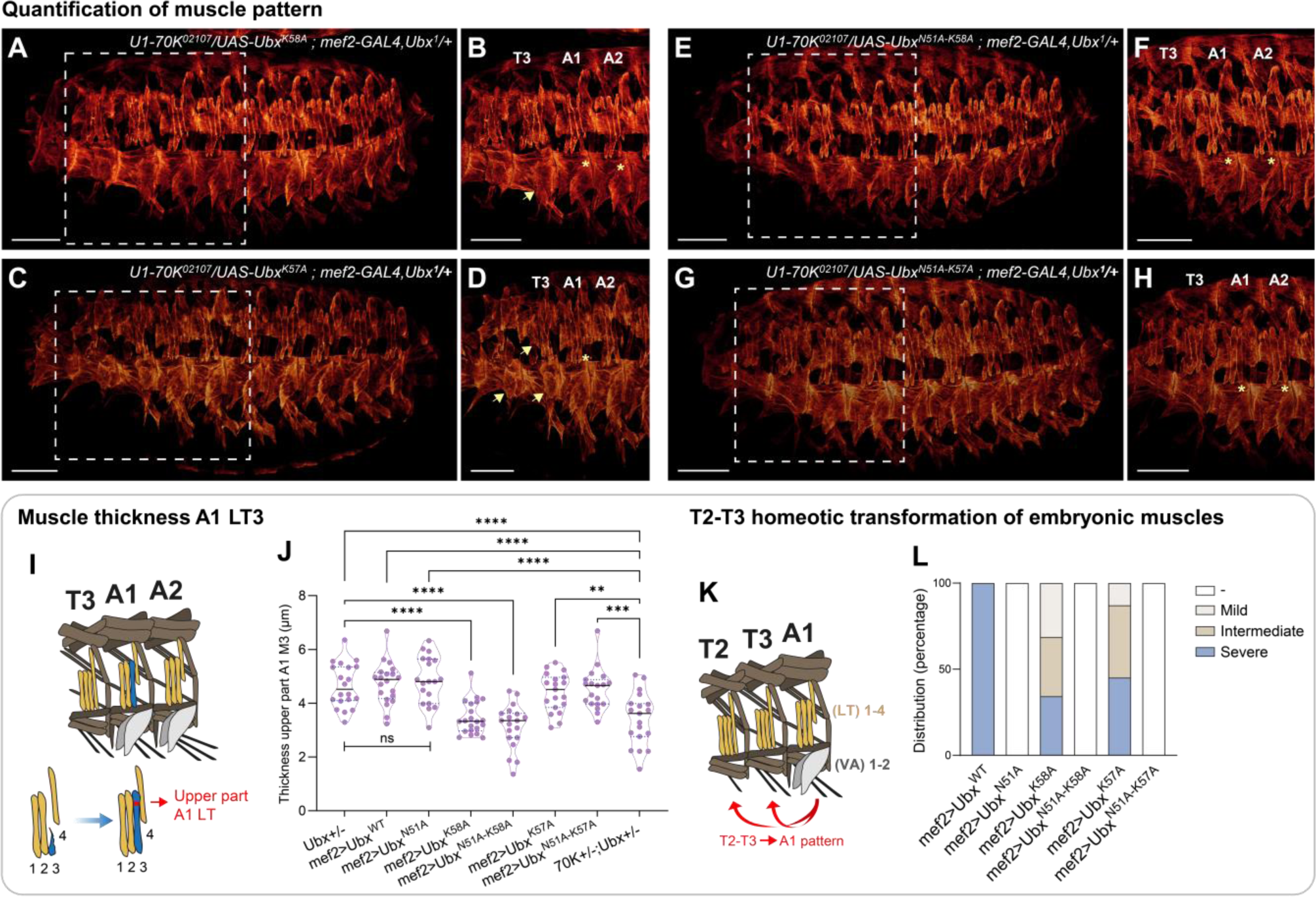
Ubx-RNA binding activity contributes to muscle morphogenesis. (**A-H**) 3D projections of stage 16 embryos and (**B, D, F, H**) zoom on T3-A2 segments (muscles stained with tropomyosin1). Rescue experiments of Ubx/U1-70K double heterozygous mutants with pan-mesodermal expression of (**A-B**) Ubx^K58A^, (**C-D**) Ubx^K57A^, (**E-F**) Ubx^N51A-K58A^ and (**G-H**) Ubx^N51A-K57A^ mutants. LT3 muscles (yellow star) and homeotic transformation (yellow arrow) are indicated. Scale bar 50 µm. (**I-J**) The muscle thickness of the A1 LT3 upper part was measured. (**I**) Muscle schematics (LT yellow, VA grey) and area quantified are indicated (red). (**J**) Violin plots representing muscle thickness with median (straight lines) and quartiles (dash lines). n=20 embryos per genotype. Statistics by one-way ANOVA (\*\**P*< 0.01, \*\*\*\**P*< 0.0001, ns = non-significant). (**K-L**) Homeotic transformation of the T2-T3 into an A1-like segment upon Ubx^WT^ expression. The phenotypes were classified into 4 categories: - = no changes (white); mild = VA muscles in T2 or T3 (beige); intermediate = VA muscles in T2 and T3 (brown); severe = VA muscles in T2 and T3 with LT muscles transformed into abdominal-like one (blue). n=20 embryos per genotype. The homeotic transformation induced by Ubx^K58A^ is milder than with Ubx^K57A^, despite a better chromatin binding ability. Ubx-RNA binding activity contributes to muscle morphogenesis and segment identity. **See also Figure S13**.

Collectively, these data demonstrated the essential *in vivo* function of Ubx-RNA binding activity in muscle morphogenesis.

### Synergistic Ubx-DNA and RNA binding activity contributes to its homeotic function

Hox TFs are fundamental coordinators of the identity and patterning of body polarity axes in metazoans. We thus investigated the homeotic potential of a Ubx-RNA binding mutant in controlling segment identity during development.

To this end, we assessed the capacity of Ubx-RNA and Ubx-DNA binding mutant transgenes to induce the homeotic transformation of thoracic T2 and T3 segments, effectively reprogramming them into abdominal-like structures. Transformation severity was classified as absent, mild, intermediate, or severe (Figure 5K-L). Mesoderm-specific expression of Ubx^WT^ resulted in severe homeotic transformation of thoracic segments (100%, Figure 1D-E). In contrast, none of the DNA binding mutants (Ubx^N51A^, Ubx^N51A-K58A^, Ubx^N51A-K57A^) induced homeotic transformation (Figure 1F-G, Figure 5E-H, 5K-L). Notably, mesoderm-specific expression of Ubx^K58A^ (RNA^mut^) and Ubx^K57A^ (DNA^+/-mut^) induced mild (M), intermediate (I), and severe (S) phenotypes, with a higher degree of homeotic transformation observed for Ubx^K57A^ (M:12.9, I:41.9, S:45.2%) compared to Ubx^K58A^ (M:31.2, I:34.4, S:34.4%) (Figure 5A-D, 5K-L). As a control, Ubx expression levels were verified (Figure S13J-Q). These findings establish that Ubx-RNA binding ability contributes to its homeotic function.

In this context, we sought to evaluate the significance of Ubx-RNA binding activity for its functional integrity. To this end, we performed rescue experiments of Ubx homozygous mutant by ectopically expressing Ubx derivatives within the endogenous Ubx expression pattern (Figure 6A-L). We employed the *Ubx^Gal4-M1^* mutant allele, which combines both the *Ubx* null mutant and GAL4 expression, mimicking endogenous Ubx expression pattern in embryos (Figure S13R). We combined *Ubx^Gal4-M1^* and *Ubx^1^* null mutant alleles with the Ubx-DNA/RNA binding mutant transgenes to evaluate their capacity to rescue the homeotic transformation of abdominal A1-A2 segments into thoracic-like identity (Figure 6A-O). We categorised the rescue degree based on LT and VA muscle phenotypes, assigning a relative score from 0 (no rescue) to 6 (full rescue) (Figure 6M-O). Rescue with Ubx^WT^ resulted in a score of 4-6. In contrast, DNA-binding mutants failed to rescue the homeotic transformation (score 0), indicating that Ubx-DNA binding activity is essential for its functional integrity. Strikingly, both Ubx^K58A^ (RNA^mut^) and Ubx^K57A^ (DNA^+/-mut^) were associated with partial rescue. However, Ubx^K58A^(RNA^mut^) exhibited a weaker rescue (score 1-4) compared to Ubx^K57A^ (score 3-4) (Figure 6M-O). This result emphasises the key contribution of Ubx-RNA binding activity in its homeotic functions. Once again, we confirmed that transgene expression levels were similar (Figure S13S-Z).

**Figure 6:**
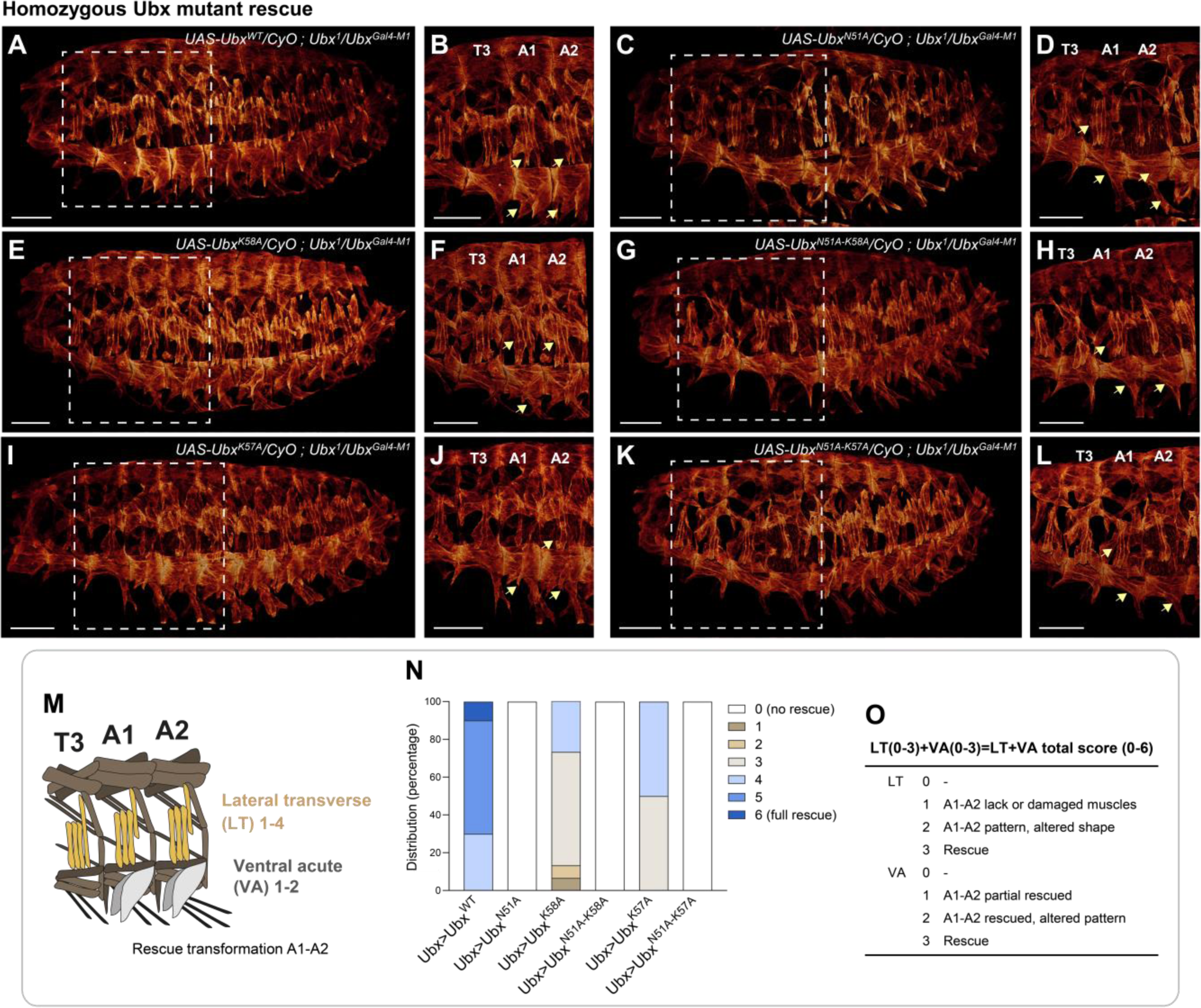
Synergistic DNA/RNA binding activity contributes to Ubx homeotic functions. (**A-L**) 3D projections of stage 16 embryos and (**B, D, F, H, J, L**) zoom on T3-A2 segments (muscles stained with tropomyosin1). Rescue experiment of Ubx homozygous mutant with Ubx-like expression (*Ubx^Gal4-M1^*) of (**A-B**) Ubx^WT^, (**C-D**) Ubx^N51A^, (**E-F**) Ubx^K58A^, (**G-H**) Ubx^N51A-K58A^, (**I-J**) Ubx^K57A^ and (**K-L**) Ubx^N51A-K57A^ mutants. Homeotic transformation and rescue of abdominal A1-A2 segments are indicated (yellow arrow). Scale bar 50 µm. (**M-O**) Distribution of homeotic transformation rescue in A1-A2 segments upon Ubx expression. (**N-O**) Phenotypes were classified according to LT and VA muscles with scores ranging from 0 (no rescue) to 6 (full rescue) (see material and methods section). n=6-14 embryos. This showed that despite a better chromatin binding ability, Ubx^K58A^ (RNA^mut^) is associated with a lower rescue of Ubx function than Ubx^K57A^ (DNA^mut^) (Chi^2^-test: p<0.0001). Thus, Ubx-RNA binding activity contributes to Ubx homeotic functions. **See also Figure S13-S14**.

Remarkably, lysine K58 is evolutionarily conserved across Hox proteins (Figure S14A). We thus extended our investigation to Hox-RNA binding conservation by evaluating the RNA binding ability of *Drosophila* Hox proteins, including the anterior Deformed (Dfd), Antennapedia (Antp), and posterior Abdominal-A (AbdA) sharing redundant functions with Ubx^42^. All Hox proteins demonstrated RNA binding potential *in vitro*. However, the K58A mutation did not affect Dfd-RNA binding and modulated Antp-RNA binding differently on distinct RNA probes, suggesting different protein-RNA interfaces across anterior and posterior Hox proteins (Figure S14B-I). Conversely, AbdA^K58A^ mirrored Ubx effects, reflecting the redundant function between the two proteins. Strikingly, while we observed a distinct behaviour of *Drosophila* Hox^K58A^, the K58A mutation of the Human Ubx-ortholog HOXA7 impaired the HOXA7-RNA binding ability similar to Ubx (Figure S14B-I). This result highlights the conservation of Hox-RNA binding ability and specificity across metazoans.

In conclusion, our results revealed that RNA binding ability is central for Ubx homeotic functions. From *Drosophila* to humans, the high conservation of Hox-RNA binding capability underlined the significance of incorporating Hox-DNA and Hox-RNA binding functions in morphogenetic networks that shape body polarity axes in all metazoans.

## DISCUSSION

Our work demonstrates novel Ubx moonlighting functions beyond its DNA binding ability. Ubx regulates differential splicing of two gene classes, regulated at transcription and splicing levels or on splicing exclusively. The latter enables the decoupling of DNA-dependent and RNA-dependent activity in splicing regulation. Notably, we identify a dynamic Ubx homodimerization-dependent mechanism shaping Ubx nuclear dynamics and Ubx-RNA association. We uncover a unique amino acid in the HD essential for Ubx-RNA binding, revealing the significance of Ubx-RNA binding for its splicing function. Our work further establishes the key role of Ubx-RNA binding for its homeotic functions in development. All in all, our findings indicate that TFs could work as integrative nodes to coordinate transcription and splicing in morphogenetic networks.

### To couple or decouple DNA and RNA dependences for Ubx splicing regulation

Our work uncovers a homodimerization-dependent mechanism for regulating alternative splicing on genes exclusively differentially spliced. Indeed, Ubx^N51A^ (DNA^mut^) retains the ability to regulate splicing and bind RNA *in vivo,* but its lack of DNA binding ability prevents its localisation near its pre-mRNA targets. In the presence of Ubx^WT^, Ubx^N51A^ is reallocated near its target transcripts and can bind RNA to exert its DNA-independent splicing function. Besides splicing factors, our results propose a unique physical connection between DNA and RNA through TF homodimerization. Our work also reveals the functional significance of this mechanism in splicing and morphogenesis.

Notably, our results indicated that the Ubx^WT^/Ubx^N51A^ cooperation on splicing is target-specific, as genes regulated both at expression and splicing levels are not differentially spliced upon Ubx^WT^/Ubx^N51A^ co-expression. Additional mechanisms promoting alternative splicing of transcripts from this gene class can be envisioned. First, Ubx could indirectly contact RNA via splicing factors, implying that Ubx^K58A^ promotes splicing. However, the K58A mutation broadly impacts Ubx splicing activity. Hence, the most plausible mechanism is a direct interaction between Ubx and RNA, which supports our result suggesting monomeric Ubx/DNA/RNA complexes (Figure S3D). Second, our previous work revealed a physical and functional interplay between Ubx and processive Pol II^21^. The Ubx/Pol II interaction could allow dynamic Ubx-chromatin loading along the gene body to efficiently couple transcription and splicing. Third, Ubx transcriptional activity implies that specific *cis*-regulatory sequences (enhancer, promoter) are recognised and bound by Ubx. Thus, the DNA sequence identity could influence the Ubx splicing function. This mechanism could involve the formation of enhancer/exon looping as proposed for VEGF or NF-κB-responsive genes^43,44^. Moreover, Hox TFs are largely enriched on promoters^21,45^. Alternatively, coupling enhancer-promoter/RNA looping and Pol II travelling could conciliate the transcription and splicing concurrent regulation (Figure 7A).

**Figure 7:**
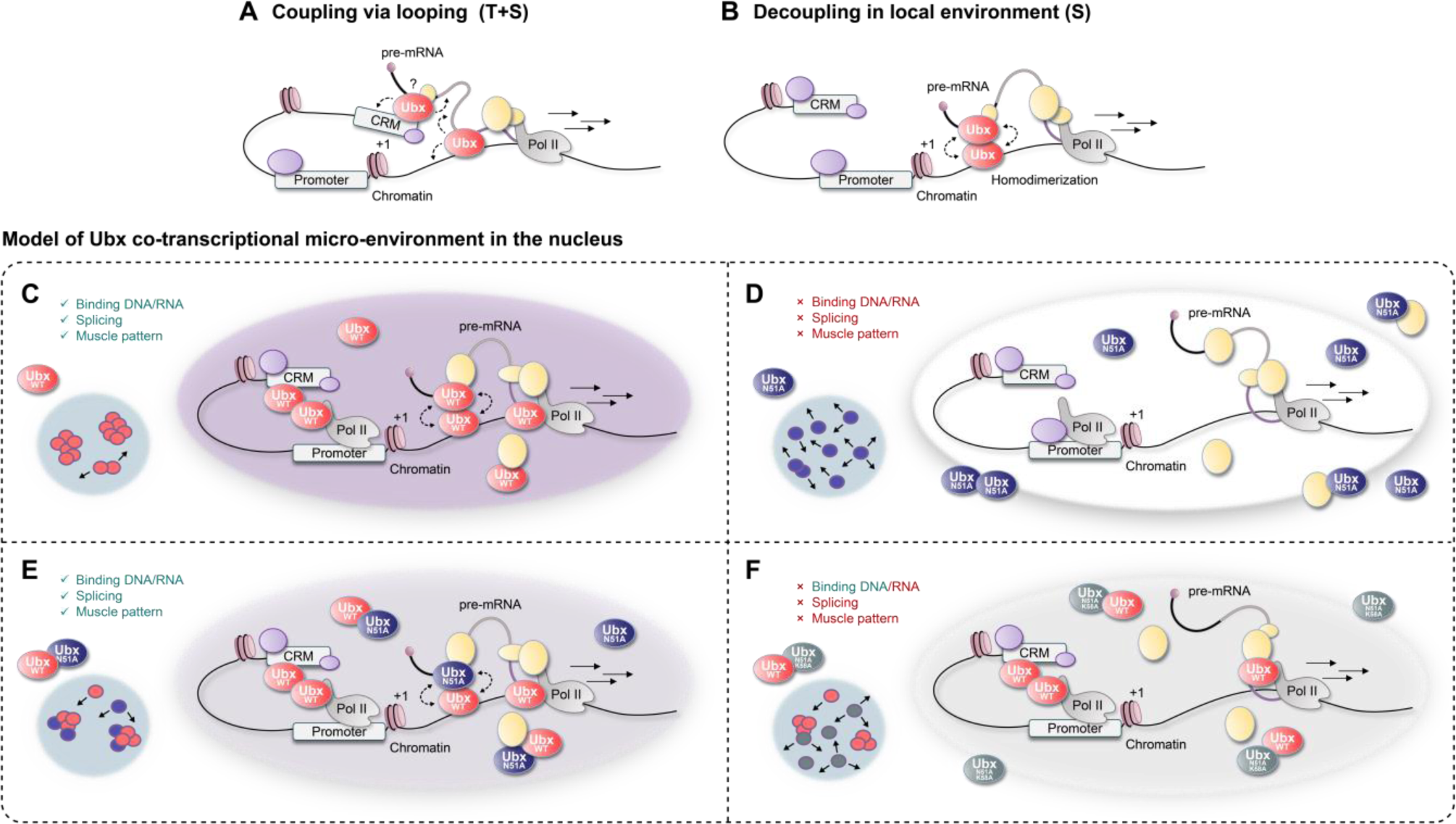
Model of Ubx co-transcriptional micro-environments. (**A-B**) Molecular model for (**A**) Ubx-target genes regulated at transcription and splicing levels (T+S) via DNA/RNA binding coupling and potentially cis-regulatory modules CRM/RNA looping. (**B**) Exclusively spliced Ubx-targets (S) are regulated by decoupled DNA and RNA binding activity with homodimerization in local environments of high TF concentration (**C-F**) Model for Ubx co-transcriptional micro-environments for regulating splicing and myogenesis. (**C**) Ubx-RNA binding ability regulates splicing globally. Ubx connects transcription and splicing functionally (T+S), or spatially (S) via a dynamic dimerization-dependent mechanism. (**D**) Ubx^N51A^ (DNA^mut^) fails to establish co-transcriptional environments. It cannot relocate near its target transcripts, thus impairing splicing and myogenesis. (**E**) The Ubx^WT^/Ubx^N51A^ dynamic dimerization facilitates the formation of co-transcriptional environments via decoupled DNA and RNA binding activity, promoting splicing and myogenesis. (**F**) Although Ubx^WT^/Ubx^N51A-K58A^ dimerize, splicing and myogenesis are impaired due to the loss of RNA binding activities.

In sharp contrast, for the Ubx-target genes solely spliced, Ubx-chromatin binding shapes protein nuclear dynamics and facilitates the formation of local environments, impacting splicing regulation via local protein concentration (Figure 7B). In this model, the target genes solely spliced could rely on a more permissive Ubx-DNA binding ability rather than sequence-specific chromatin binding. Besides, Tsai and colleagues found that Ubx forms nuclear micro-environments that are promoted by low-affinity DNA binding sites, suggesting that local TF concentrations help these sites overcome inefficiency^46^. Given that Ubx-DNA binding is essential for its splicing activity, it will be interesting to investigate how DNA binding affinity influences Ubx splicing activity on the various gene classes differentially spliced.

### Model of Ubx co-transcriptional micro-environment

Our study demonstrated that Ubx forms functional and dynamic dimers crucial for splicing regulation by decoupling its DNA and RNA binding functions. Interestingly, many TFs dimerize to modulate DNA binding specificity, which seems to be an evolutionarily conserved feature^47^. Various data suggest that HD-TF dimerization affects their DNA binding function^38,48,49^. In addition to transcription initiation, transient TF homodimerization could aid the maintenance of local co-transcriptional environments. Ubx micro-environments are characterised by high Ubx protein concentrations and associated with active transcription^46^. Beyond transcription per se, local TF concentration and binding sites affinity might be crucial for fine-tuning co-transcriptional regulation, favouring RNA binding and Ubx-splicing activity or DNA binding and transcription. Combined with our results, we proposed a refined model in which Ubx forms co-transcriptional micro-environments wherein transcription and splicing processes are spatially connected through either functional coupling (*Chas*) or decoupling (*Pura*) on distinct target genes (Figure 7C-F). We proposed that these co-transcriptional micro-environments are essential to facilitate the orchestration of multilayered gene regulatory networks by TFs.

Moreover, specific splicing regulation likely depends on distinct protein networks between Ubx and splicing factors, which could fine-tune Ubx-specificity and affinity toward its target transcripts. Ubx could act as an interactive platform to recruit specific splicing factors for fine-tuning alternative splicing. However, it is still unclear whether these protein complexes are formed primarily on the nucleoplasm, the chromatin, or nascent transcripts. Considering the diversity in TF-dimerization interfaces, local TF concentration coupled with DNA/RNA binding specificity and TF-splicing factor interaction could fine-tune the TF-specificity on transcription and/or splicing on target genes in local co-transcriptional micro-environment.

### K58A impact and Hox-RNA binding activity in morphogenesis

Our work identified and characterised the Ubx^K58A^ RNA binding mutant, representing a breakthrough in studying Ubx function. This enabled us to decipher the dual role of Ubx as a transcription factor and splicing regulator, demonstrating the contribution of its RNA binding activity to splicing and muscle morphogenesis (Figure 7F). Although numerous factors likely fine-tune specificity or affinity, our comprehensive analyses, from *in vitro* to *in vivo* experiments, underscore the importance of the HD lysine K58 for RNA binding and the contribution of Ubx-RNA binding activity for splicing and muscle development. Notably, we uncovered a dimerization-dependent mechanism for Ubx to regulate splicing and highlighted its critical role in morphogenesis. In future, it would be interesting to explore whether similar interactions with other splicing factors can be functionally rescued, revealing a potential specificity of this mechanism. Moreover, we demonstrated that Ubx-RNA binding activity is crucial for muscle morphogenesis and segment identity, independently from its transcriptional effects. This is supported by our results showing that Ubx^K57A^ (DNA^+/-mut^) has a more significant impact on chromatin binding than the K58A (RNA^mut^) mutation and still depicted a better rescue of segment identity (Figure 6). Thus, our results establish the key role of Ubx-RNA binding activity and potentially splicing in segment identity. As RNA-binding is conserved across the Hox family (Figure S14), further investigation in various tissues and with other Hox proteins will help to generalise the importance of Hox-RNA binding activity for their homeotic functions.

Hox TFs are crucial for the development of body plans in metazoans. Their function has been extensively described at the transcriptional level. Beyond this conventional function, our data demonstrated the importance of their RNA-binding function for splicing and morphogenesis. Our work further emphasises that incorporating their DNA and RNA binding functions offers a more comprehensive view of their regulatory roles *in vivo*. In future, we believe that this inclusive approach to TF molecular functions will lead to novel insights into developmental biology, evolution and potential biomedical applications.

## ACKNOWLEDGEMENTS

Stocks obtained from the Bloomington Drosophila Stock Center (NIH P40OD018537) were used in this study. The BB5/37.1-s antibody developed by Belinda Bullard was obtained from the Developmental Studies Hybridoma Bank, created by the NICHD of the NIH and maintained at The University of Iowa (Department of Biology, Iowa City, IA 52242). We thank the FlyORF injection service (Zurich, Switzerland) for the transgenic fly line generations. We warmly thank Ingrid Lohmann for sharing the homemade antibody Ubx guinea pig. We thank the PLATIM imaging facility and especially Elodie Chatre for her support in setting up the FRAP experiments and the IGFL for the unrestricted access to the imaging facility and hosting. We thank Pedro Pinto for the discussions on the project and the introduction to 3D microscopy acquisition. We are very grateful to Nawal Hajj-Sleiman, Pedro Pinto, Didier Auboeuf, Mounia Lagha and Ingrid Lohmann for insightful discussions, their feedback and advice on the manuscript.

## AUTHOR CONTRIBUTIONS

Conceptualisation: JC. Experimental Design: JC, CB. Experimental procedures: CB, WX, PB, MS, ASH, JC. Formal analysis: JC, WX. Supervision: JW, JC. Resources: SM. Data Visualisation: PB, JC. Writing and editing: JC with the support of all authors. Project administration: JC. Funding acquisition: JC.

## SUPPLEMENTAL INFORMATION

Document S1. Figure S1-S14

The resource table is included in the method section at the end of the manuscript

## MATERIALS AND METHODS

### Resources Table

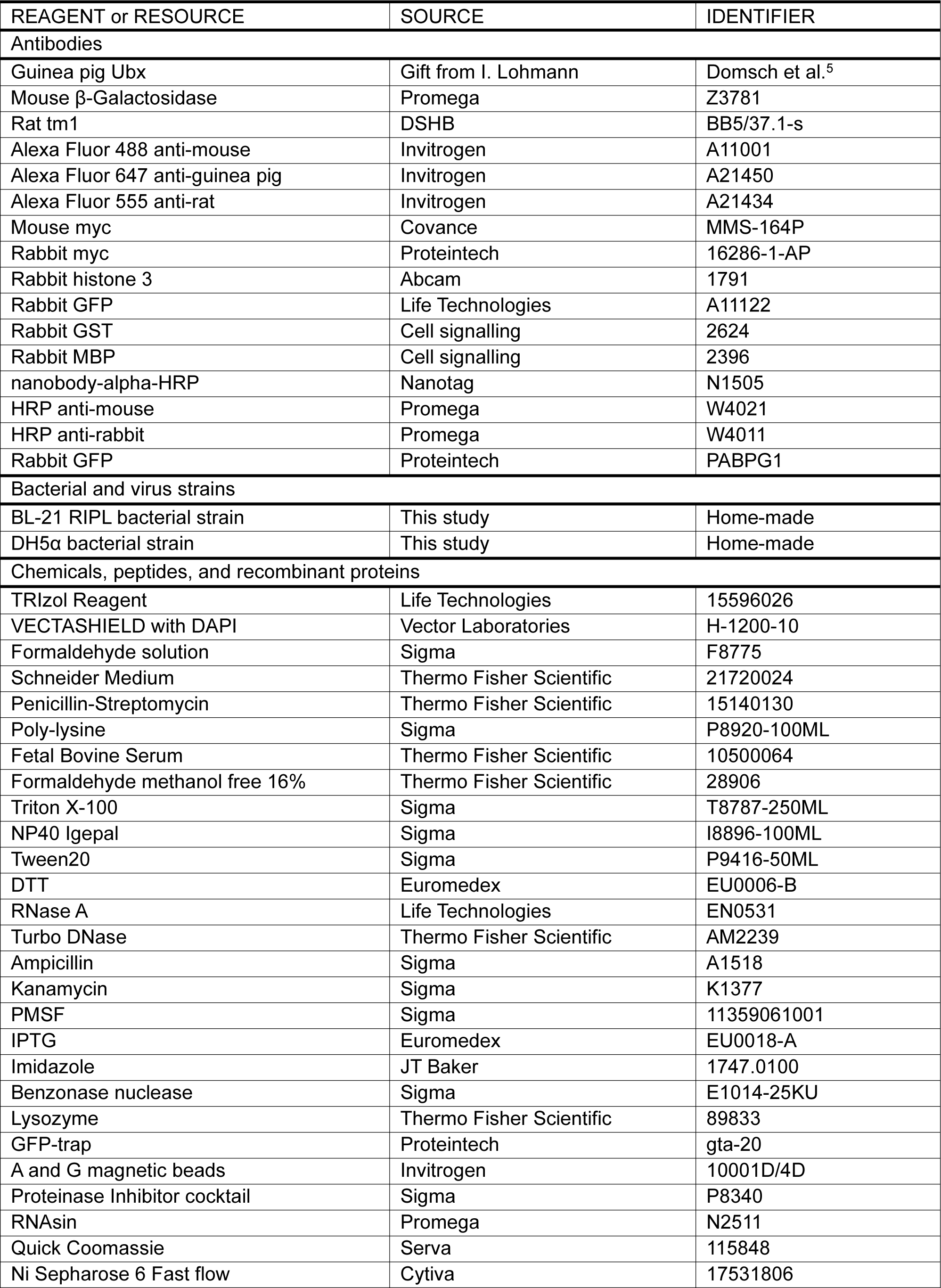
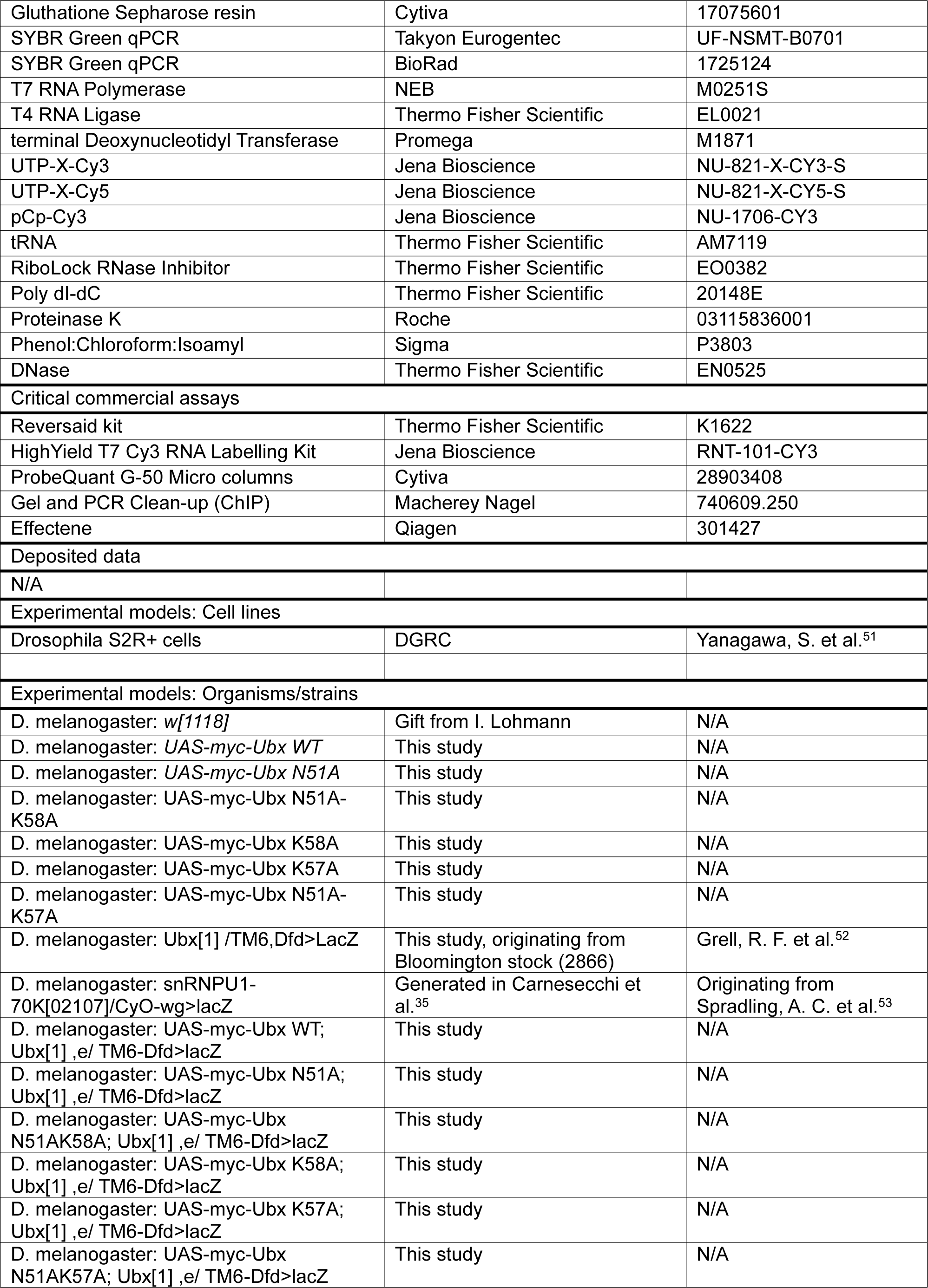
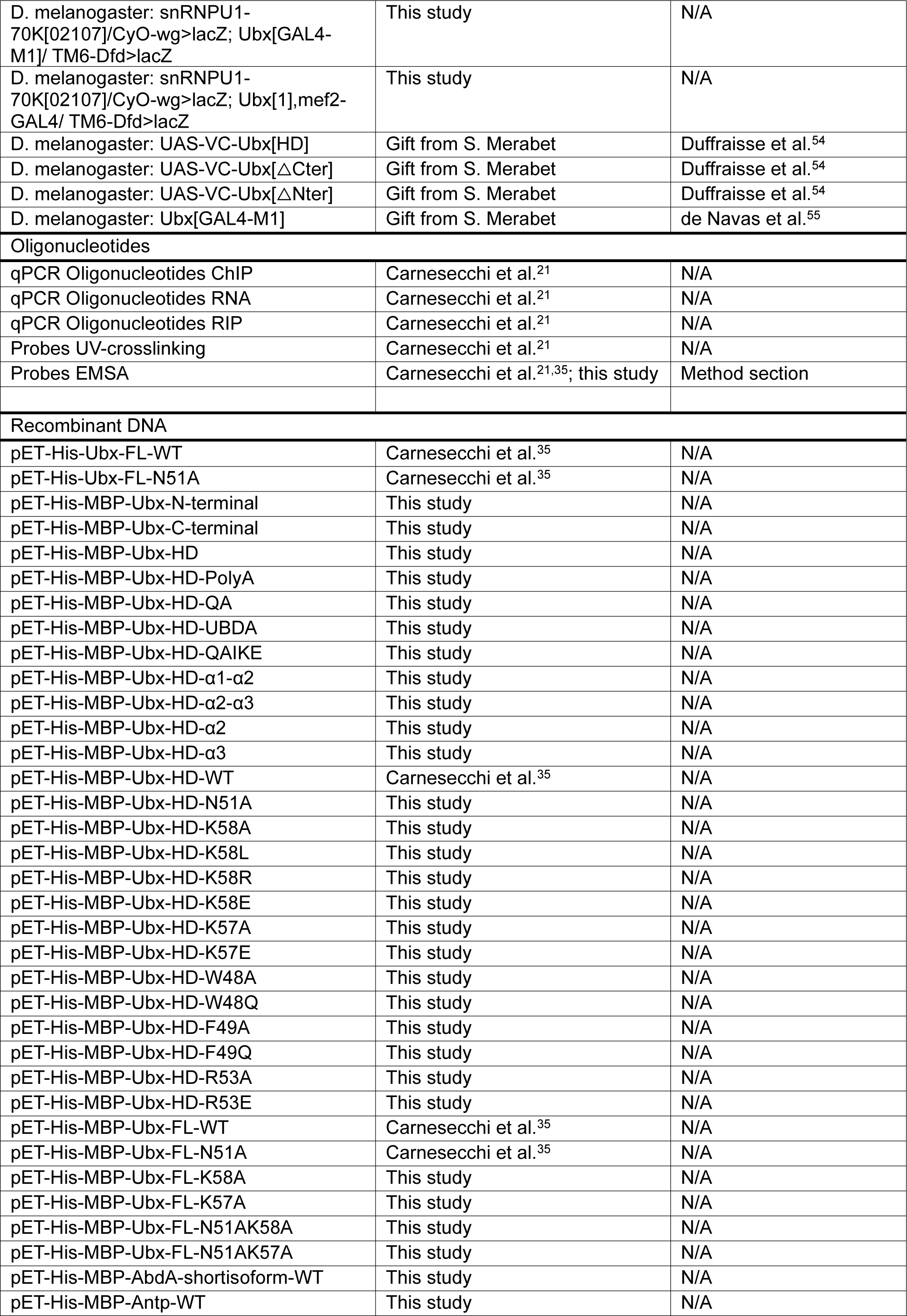
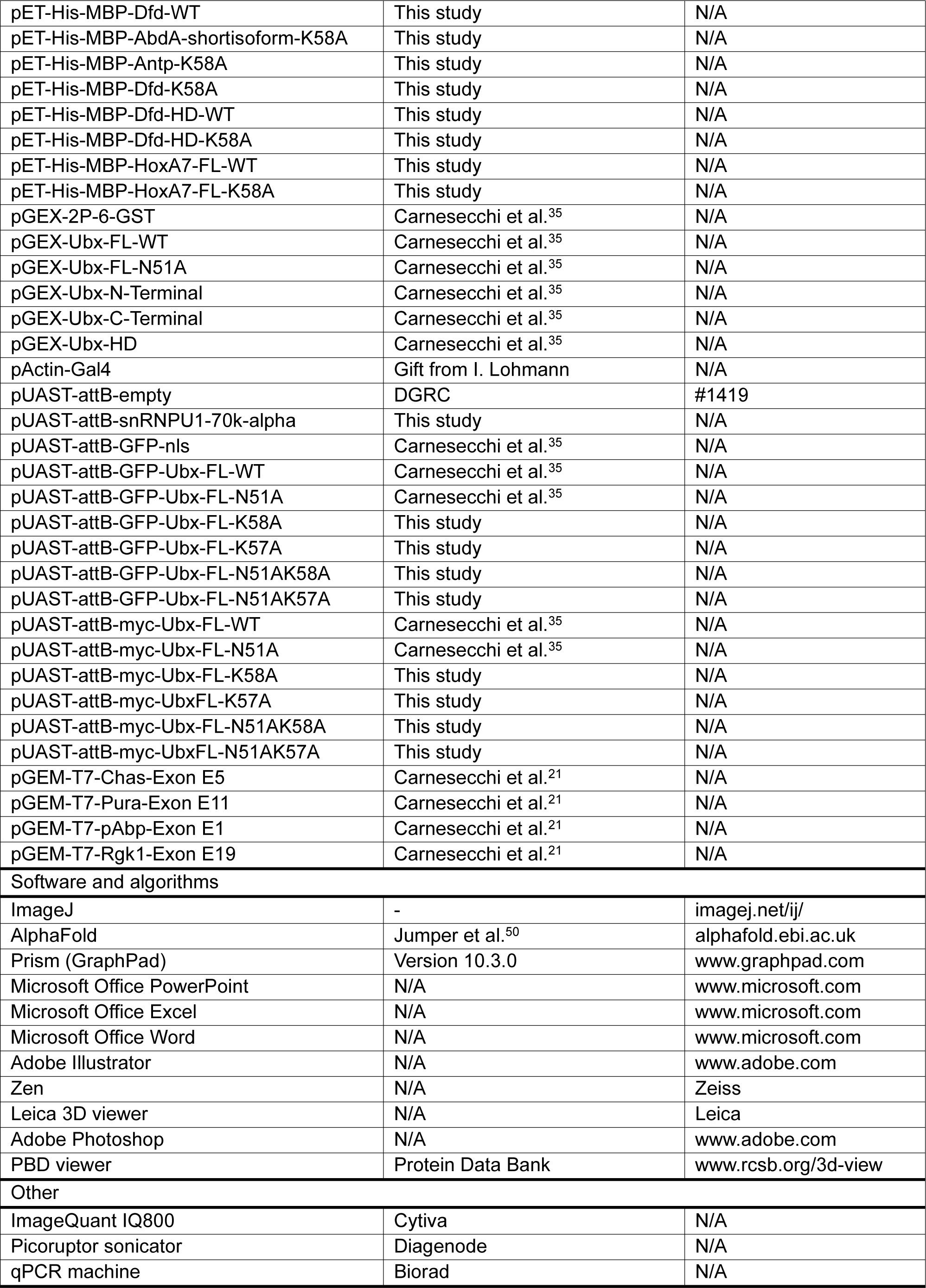

### Materials availability

Plasmids and *Drosophila* transgenic lines generated in this study will be available upon reasonable request from the lead contact, Julie Carnesecchi.

### Biological Resources

*E.coli* DH5α cells were used to produce and purify plasmids. *E.coli* BL-21 RIPL were used to express and purify recombinant proteins. The fly lines used for the study are listed and documented in the resource table. Transgenic lines were generated by φC31-mediated transgenesis on the VK37 landing site^56^ (chromosome 2) and listed in the resource table. *Drosophila* S2R+ cells were obtained from the *Drosophila* Genomic Research Centre (DGRC) and maintained at 25°C. The vector constructs used in the study are listed and documented in the resource table. Primers are listed in ^21^ without any changes. Probes are listed in ^21^ and additional probes are included in the method section.

### Immunofluorescence and imaging

14-17 staged embryos were collected after 5 hours of laying with an additional 12 hours of ageing at 25°C. Embryos were dechorionated with 100% bleach for 3 minutes, washed with water, and fixed with formaldehyde supplemented with heptane. The vitelline membrane was removed using methanol. The devitellinised embryos were collected and washed two times with methanol. For immunostaining, embryos were washed in PBS-Tween 0.1% for 10 minutes three times, blocked with BSA 1% in PBS-Tween for one hour, and incubated with primary antibodies overnight at 4°C. Secondary antibodies coupled to a fluorescent protein (Life Technologies, 1/300e) were incubated for 2 hours the following day, and embryos were mounted in Vectashield-DAPI (Vectorlabs) to stain nuclei. The following antibodies were used: Ubx (1/500e, Home-made), Beta-Galactosidase (1/1000e, Promega, Z3781), tropomyosin1 tm1 (1/200e, DSHB, BB5/37.1-s). Images were acquired on the Leica SP8 confocal microscope using a standard plan-Apochromat 40x, NA 1.3, Oil objective. The collected images were analysed and processed using the Leica program for 3-dimensional (3D) views and Fiji. For quantification, all images were taken with unique parameters.

#### Muscle thickness quantification

tm1 staining was used to visualise the muscle pattern and Beta-Galactosidase to distinguish embryo genotypes. The imaging settings included: scan speed 600 Hz; resolution 1024x1024 pixels; Z-stack of 25-45 sections (step size 0.8μm) to capture a 3D structure of embryonic hemi-segment. Images were then analysed by Fiji software. The scale was identical for all quantified pictures. The four lateral transverse muscle thicknesses of A1-A2 segments were measured for at least 20 embryos per genotype.

#### Ubx level quantification

A stack of 10 z-slices (step size 0.8μm) covering the Ubx signal was selected to quantify Ubx and DAPI intensities in each embryo. For each z-slice, a relative signal was obtained by calculating the ratio intensity (mean gray value) for Ubx signal (488 nm) relative to the DAPI signal in T2-T3 segments (which only expressed ectopic Ubx protein) using the same region of interest (ROI). The mean signal ratio for each stack (or embryo) was calculated and plotted. Quantification was performed for 5-15 embryos per genotype.

#### Homeotic transformation score

For Ubx/U1-70K heterozygous mutant with T2-T3 transformation, the transformation was scored as follows for 20 embryos: -: no changes, mild: VA muscles visible in T2 or T3, intermediate: VA muscles visible in T2 and T3, severe: VA muscles visible in T2 and T3 with LT muscles transformed into abdominal like phenotype.

For Ubx homozygous mutant rescue experiments, the transformation rescue was scored according to VA and LT identities for 5-13 embryos. A score of 0-3 was used for VA and LT, and the sum was plotted (0 to 6 for complete rescue). The score was set as follows: for LT muscles, “0” represents homeotic transformation (A1-A2→T3, no rescue); “1” represents A1 or A2 segments with 1-2 muscles missing with alteration; “2” indicates LT muscles showing the A1 abdominal pattern with no muscle missing but alteration; “3” indicates A1-A2 rescue, with muscle patterns similar to the control. For VA muscle transformation, the scoring was: “0” for homeotic transformation; “1” for partial rescue of A1 or A2 VA muscles; “2” for A1 and A2 VA muscles rescued but with damaged muscle shapes; “3” indicates A1 and A2 rescue.

### Cell culture and transfection

*Drosophila* S2R+ cells were maintained at 25°C in Schneider medium supplemented with 10% FCS, 10 U/ml penicillin and 10 µg/ml streptomycin. Cells were simultaneously seeded and transfected with Effectene (Qiagen) according to the manufacturer’s protocol. The GAL4/UAS system was used for inducible protein expression driven by the Actin promoter (pActin-GAL4). For RNA analysis, cells were seeded in 6 well plates and transfected as described with UAS-GFPnls or -myc-Ubx constructs. For interaction, ChIP and RIP assays, 10.10^6^ cells were seeded in 100 mm dishes and transfected as described with UAS-GFPnls, UAS-GFP-Ubx combined with UAS-myc-Ubx constructs, and pActin-GAL4. Cells were harvested in PBS after 48 hours of transfection, and pellets were resuspended with lysis buffer supplemented with protease inhibitor cocktail (Sigma), 0.1 mM PMSF and 1 mM DTT. For FRAP analysis, cells were seeded and transfected (50 ng of each plasmid) in 12 well plates and transferred with fresh supplemented media in glass bottom dishes coated with Poly-lysine (Sigma) at least 2 hours before image acquisition.

### RNA extraction, retrotranscription (RT) and quantitative PCR

Total RNAs for 3 to 5 independent experiments performed in triplicate were extracted by Trizol (Life Technologies) and quantified on NanoDrop. The samples were digested with DNase (Thermo Fisher Scientific), and 1 μg of RNA was converted to first strand cDNA using Reversaid kit (Thermo Fisher Scientific) and random hexamers in 20 µl final volume, according to the manufacturer’s protocol. Real-time PCR experiments have been performed according to MIQE guidelines ^57^ in CFX96 Real-time systems (BioRad). In detail, qPCR was performed in technical duplicate for each sample, in a 96-well plate using the SYBR™ Green (BioRad and Takyon) in a final volume of 10 µl. Primers (exactly the same as listed in ^21^) were designed with Primer3 for amplicons ranging between 70-150bp, verified with nucleotide blast (NCBI) and tested by serial dilution of cDNA and melt curve analysis. qPCR cycles were: 95°C for 2 minutes, 40 cycles of: 95°C, 15 seconds; 60°C, 30 seconds, followed by temperature gradient. Data were quantified by the ΔΔ-Ct method and normalised to Actin 5C expression or internal region of constitutively expressed exons of the related gene as indicated.

### Nuclear RNA-immunoprecipitation and quantitative PCR

Confluent *Drosophila* S2R+ cells plated in 100 mm dishes were collected in cold PBS 48 hours after transfection, as in ^21^. After several PBS washes, cell pellets were resuspended in buffer A (10 mM Hepes pH 7.9, 10 mM KCl, 1.5 mM MgCl2, 0.34 M sucrose, 10% glycerol). Lysates were incubated with 0.1% Triton for 5 minutes and centrifugated at 1300 rcf. Nuclear pellets were resuspended with IP buffer (150 mM NaCl, 50 mM Tris pH 7.5, 1 mM EDTA, 1% Triton), incubated on ice with vigorous regular vortex and sonicated (2x 30 seconds on/off, Picoruptor, Diagenode). All buffers were supplemented with a protease inhibitor cocktail (Sigma), 1 mM DTT, 0.1 mM PMSF and RNasin (Promega). Input fractions were collected both for protein and RT-qPCR control. 1-1.5 mg of nuclear lysates were diluted in IP buffer, pre-cleared with 15 µl A/G plus agarose beads (Santa Cruz) and incubated for 5 hours with 20 µl of GFP-Trap beads (Proteintech). Beads were washed 5 times for 5 minutes at 4°C with rotation and IP buffer. 10% were collected for protein analysis, the remaining beads were resuspended in Trizol (Life Technologies), and RNAs were extracted according to the manufacturer’s protocol. Notably, RNAs were precipitated overnight at -70°C in isopropanol to increase yield. Retro-transcriptions were processed, as described in the method section, on 2 µg of RNA by doubling the total reaction volume (40 µl final). 20 µl of water was added to cDNAs, and 2 µl were used for quantitative PCR. Enrichment was calculated relative to input, which referred to the total RNA present within each sample and GFP control values. Sequences of the primers used in this study are provided in ^21^.

### Chromatin-immunoprecipitation coupled with quantitative PCR

48 hours post-transfection, confluent *Drosophila* S2R+ cells plated in 100 mm dishes were crosslinked with 1% formaldehyde and quenched for 5 minutes in 0.125 M Glycine as described in ^21^. After several PBS washes, cell pellets were resuspended in lysis buffer (1% SDS, 50 mM Tris·HCl, pH 8, 10 mM EDTA). Sonication (2 cycles, 30 seconds on/off) was performed with Picoruptor (Diagenode) after verification of the fragmentation profile (bioanalyser). After centrifugation, input samples were taken, and the remaining lysates were diluted in the dilution buffer (0.01% SDS, 1% triton, 2 mM EDTA, 20 mM tris pH 8,1, 150 mM NaCl). The diluted lysates were incubated with 2 μl of GFP antibody (Proteintech, PABG1) overnight at 4°C on rotation and for two additional hours with mixed Dynabeads protein G and A (15:15 µl, Life Technologies). Beads were washed with TSE-150 (0.1% SDS, 1% Triton, 2 mM EDTA, 20 mM Tris, pH 8.1, 150 mM NaCl), TSE-500 (as TSE-150 with 500 mM NaCl), LiCl detergent (0.25 M LiCl, 1% NP40, 1% Sodium Deoxycholate, 1 mM EDTA, 10 mM Tris, pH 8.1), and twice with Tris-EDTA (5:1 mM). Combined elution and decrosslinking were performed by adding RNase A for 30 minutes at 37°C, then 0.1% SDS with proteinase K for 1 hour at 37°C and additional incubation with NaCl under 900 rpm shaking for 7 hours at 65°C. DNA fragments were purified using Qiaquick miniElute (Qiagen) according to the manufacturer protocol and diluted to 1/10 for input and 1/2 for immunoprecipitated fractions. qPCRs were performed using 2 μl of DNA, and enrichment was calculated relative to input and Ubx^N51A^ derivatives values. The immunoblotting represents 2% of the cell lysate used for the IP. Sequences of the primers used in this study are provided in ^21^.

### Co-immunoprecipitation of whole cell lysate

For co-immunoprecipitation assays, transfected *Drosophila* S2R+ cells were harvested in PBS, and the pellets were rinsed with PBS. Pellets were resuspended in NP40 buffer (20 mM Tris pH 7.5, 150 mM NaCl, 2 mM EDTA, 1% NP40) and treated with Benzonase (Sigma) as described in ^21^. For crosslink, *Drosophila* S2R+ cells were crosslinked with 1% formaldehyde and quenched for 5 minutes in 0.125 M Glycine. After several PBS washes, cell pellets were resuspended in NP40 buffer, sonicated (3 cycles, 30 seconds on/off, Picoruptor, Diagenode) and treated with Benzonase. GFP-Trap beads (Proteintech) were added to 1-1.5 mg of protein extract, incubated for 2 hours and washed 5 times with NP40 buffer with vortex. Input fractions represent 1-10% of the immunoprecipitated fraction.

### SDS-PAGE and Immunoblotting

For western blot analysis, proteins were resolved on 8-15% SDS-PAGE, blotted onto PVDF membrane (Millipore) and probed with specific antibodies after saturation. The antibodies (and their dilution) used in this study were: myc (mouse Covance, rabbit Proteintech, 16286-1-AP 1/3000e), histone 3 (Abcam, 1791, 1/20,000e), GFP (Life Technologies, A11122, 1/3,000e), GST (Cell signalling, 2624, 1/5,000e), MBP (Cell signalling, 2396, 1/3,000e), nanobody-alpha-HRP (Nanotag, N1505, 1/2,000e)

### Protein purification and GST pull-down

His-tagged and GST-tagged proteins were cloned for this study or from our previous work ^21,35^ in pET or pGEX-6P plasmids, respectively, and are listed in the resource table. His- and GST-tagged proteins were produced from BL-21 (RIPL) bacterial strain, purified on Ni-NTA agarose or Glutathione-Sepharose beads (GE-healthcare), respectively and quantified by Coomassie staining. His-tagged proteins were specifically eluted from the beads with Imidazole. *In vitro* interaction assays were performed with equal amounts of GST or GST fusion proteins in affinity buffer (20 mM Hepes, 10 µM ZnCl_2_, 0.1% Triton, 2 mM EDTA) supplemented with NaCl, 1 mM DTT, 0.1 mM PMSF and protease inhibitor cocktail (Sigma). Proteins produced *in vitro* were subjected to interaction assays for 2 hours at 4°C under mild rotation with *Drosophila* nuclear extracts or recombinant protein produced and purified *in vitro*. Bound proteins were washed 4 times and resuspended in Laemmli buffer for western-blot analysis. The input fraction was loaded as indicated.

### *In vitro* transcription with Cy3-UTP labelling

For *in vitro* transcription, we employed selected fragments of alternatively spliced exons that were cloned into the pGEM®-T Easy (Promega) Vector (listed in ^21^). To generate the DNA templates for transcription, plasmids were amplified in DH5α bacterial strain, purified and linearised 3’ to the cloned sequence using the SpeI restriction site. Internally labelled RNAs were produced by *in vitro* transcription using the HighYield T7 Cy3 RNA Labelling Kit (Jena Bioscience, RNT-101-CY3) following the manufacturer’s instructions. Each reaction contained 500 ng DNA template, 0.4 μl Cy3-UTP (5 mM) and 0.4 μl RiboLock RNase Inhibitor (Thermo Fisher Scientific) and was incubated for 2 hours at 37°C. DNA template was digested with 1 μl TURBO™ DNase (Thermo Fisher Scientific) for 15 minutes at 37°C. Finally, labelled RNA probes were purified using the ProbeQuant™ G-50 Micro Columns (Cytiva) and eluted in 50 μl.

### *In vitro* transcription with 3’-Cy3 labelling

Cold *in vitro* transcription was performed in a volume of 50 μl containing 1 μg DNA template (annealed oligonucleotides), 1 mM NTPs solution, 100 U RiboLock RNase Inhibitor (Thermo Fisher Scientific), 1x Transcription Buffer and 500 U T7 RNA Polymerase (NEB). Following incubation of the reaction for 2 hours at 37°C, the DNA template was digested with 2.5 μl TURBO™ DNase (Thermo Fisher Scientific) for 15 minutes at 37°C. Unlabelled RNA was purified using the ProbeQuant™ G-50 Micro Columns (Cytiva) and eluted in 50 μl. To concentrate the produced RNA, precipitation was carried out with 0.3 M sodium acetate pH 5.2 and 2.5 volumes of ethanol 100%, incubation for 30 minutes at - 80°C and centrifugation at maximum speed for 30 minutes at 4°C. After air-drying the pellet for 5 minutes, RNA was resuspended in 15 μl DEPC-treated water. Finally, the 3’-Cy3 labelling reaction was performed overnight at 4°C in 20 μl containing 100 pmol unlabeled RNA, 0.5 mM ATP, 200 pmol pCp-Cy3 (Jena Bioscience), 10 U RiboLock RNase Inhibitor (Thermo Fisher Scientific), 1x Ligation Buffer and 10 U T4 RNA Ligase (Thermo Fisher Scientific). Labelled RNA probes were purified using the ProbeQuant™ G-50 Micro Columns (Cytiva) and eluted in 50 μl.

### Protein-RNA UV-crosslinking assay

To prepare the protein-RNA complexes for UV-crosslink, 2 pmol of internally labelled RNA probes were mixed with approximately 0.5-1 μg of his- or his-MBP-purified proteins. The binding reaction was performed in a pre-cooled 96 well plate in a volume of 30 μl containing 1x binding buffer (20 mM Hepes pH 7.9, 1.4 mM MgCl2, 1 mM ZnSO4, 40 mM KCl, 0.1 mM EDTA, 5% Glycerol), 2 μg tRNA (Thermo Fisher Scientific), 3 μg BSA, 10 mM DTT and 0.1% NP40. After 20 minutes on ice, the samples were irradiated with UV light in a UVP-Crosslinker (Jena Analytik) for 10 minutes on ice and subsequently transferred to Eppendorf Tubes. 1.5 μl of RNase A (Life Technologies) were added and the samples were incubated for 20 minutes at 37°C. Cy3-labelled protein-RNA complexes were resolved on 10-12% SDS-PAGE for 40 minutes at 180 V and detected by fluorescence using Imager (IQ800, Cytiva). Following the detection, the gels were stained with Coomassie overnight, rinsed with water and imaged.

### Electrophoretic Mobility Shift Assay (EMSA)

5’Cy5-DNA probes were produced commercially and 3’-UTP-Cy5 ligation was performed at 37°C on double-strand DNA with terminal Deoxynucleotidyl Transferase (TdT) (Promega). The 5’-Cy5 labelled complementary oligonucleotides were annealed before the reaction. The sequences are Ubx/Exd site: TTCAGAGCGAATGATTTATGACCGGTCAAG, Chas exon 5:TAATCAATAGCCAAAGAGCTA-CTGCTGCTGCTGCTGCTGCTGCTGCTCCTGCTTGGCTa. The binding reaction was performed for 20 minutes in a volume of 30 μl containing 1x binding buffer (20 mM Hepes pH 7.9, 1.4 mM MgCl2, 1 mM ZnSO4, 40 mM KCl, 0.1 mM EDTA, 5% Glycerol), 0.2 μg Poly(dI-dC) for DNA EMSA, 2 µg of tRNA for RNA EMSA, 0.1 μg BSA, 10 mM DTT and 0.1% NP40. For RNA EMSA, the reaction mix was heated for 3 minutes at 70°C and immediately placed on ice for 3 minutes. For each reaction his-purified proteins were used. Separation was carried out (1 hour, 150 V) at 4°C on a 4-6% acrylamide gel in 0.5x Tris-borate-EDTA buffer to visualise complex formation by retardation. Cy5-labelled DNA-protein and Cy3-labelled RNA-protein complexes were detected using IQ800 Cytiva Imager.

### Fluorescence Recovery After Photobleaching acquisition and modelling

Time series were acquired on a confocal Zeiss LSM710, Axio Observer Z1 equipped with a plan-apochromat 63x, NA 1.4, Oil objective and confocal Zeiss LSM780 with plan-apochromat 63x, NA 1.4, Oil objective, both equipped with FRAP module. Spectral detection was done on GaAsP 32+2 PMT, with a conventional scanner. Half nuclei were bleached to reach 50% fluorescence drop at 100% laser power (0,45 mW). Duration for one frame was 614.4 milliseconds for 80 cycles including 5 pre-bleach frames, for a total duration of 52 seconds which mirrors the same profile model in ^21^ over 4.5 minutes, thus avoiding cell drift over long acquisition. Acquired time series were analysed using Zen software with the integrated FRAP module. The recovery curves were fitted with the double exponential diffusion models (I= IE-I1 x exp (-t/T1) – I2 x exp (-t/T2)) as previously validated for Ubx molecules (^21^validated model with Akaike and Bayesian information criterion). The area of interest (ROI) was normalised to the background and reference ROI in the non-bleached nuclei region. The fitted data of the recovery curve were visualised with GraphPad Prism. The t-half number and population fractions were calculated with the integrated module of the Zeiss Zen software for 25-44 nuclei per condition.

### Data analyses and visualisation

Data visualisation was achieved with Fiji (is just ImageJ), GraphPad Prism 10, Microsoft Office PowerPoint, Excel and Adobe Illustrator and Photoshop software. Gel and phenotype quantifications were performed with Fiji (is just ImageJ). Statistical analyses were performed using one-way ANOVA, Chi^2^ for distribution and t-test multiparametric using GraphPad Prism 10 software. Each experiment was performed for 3 to 5 independent biological replicates for the gels and immunoblots, additional technical duplicates for the RIP-qPCR and ChIP-qPCR and technical triplicates for RNA expression. Final calculations, tests for significance and graphic illustrations were performed with GraphPad Prism 10.

