## Supplemental files for "Synergistic DNA and RNA binding of the Hox transcription factor Ultrabithorax coordinates splicing and shapes *in vivo* homeotic functions"

### SUPPLEMENTAL INFORMATION

**Figure S1-S14**

#### Synergistic DNA and RNA binding of the Hox transcription factor Ultrabithorax coordinates splicing and shapes *in vivo* homeotic functions


**AUTHORS**

Constanza Blanco[1#], Wan Xiang[1,2,3#], Panagiotis Boumpas[1,4], Maily Scorcelletti[1,4], Ashley Suraj Hermon[1], Jiemin Wong[2], Samir Merabet[1], Julie Carnesecchi[1,3,5*]

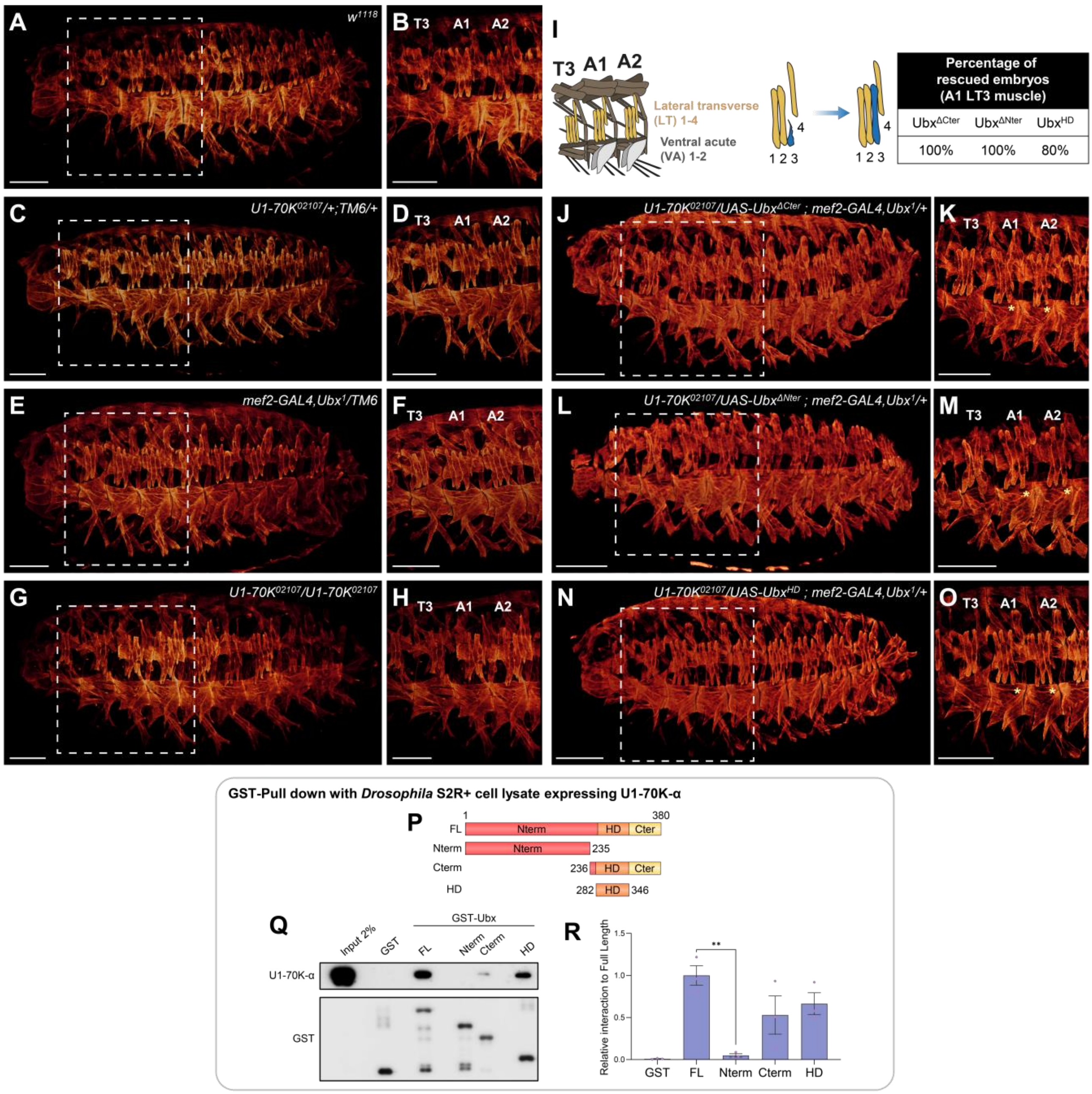


**Figure S1: U1-70K and Ubx HD genetically and physically interact *in vivo* and *in vitro*.** (**A-H, J-O**) 3D projections of stage 16 embryos (muscles stained with tropomyosin1) and (**B, D, F, H, K, M, O**) zoom over T3-A2 segments for (**A-B**) control $w^{1118}$, (**C-D**) U1-70K heterozygous mutant, (**E-F**) Ubx heterozygous mutant, (**G-H**) U1-70K homozygous mutant, associated with a global alteration of the LT muscles in all segments. n=20 embryos for control and heterozygous, n=5 embryos for homozygous mutant embryos. (**I**) Cartoon of muscle pattern and rescue percentage for Ubx derivatives expressed in mesoderm in Ubx/U1-70K genetic interaction background. Rescue experiments of double heterozygous (*snRNP*)$U1\text{-}70K^{02107}$ with $Ubx^{1}$ embryos with mesoderm-specific expression (*mef2-GAL4*) of Ubx derivatives, (**J-K**) deleted from the C-terminal ($UAS\text{-}Ubx^{\Delta Cter}$), (**L-M**) deleted from the N-terminal domain ($UAS\text{-}Ubx^{\Delta Nter}$), or (**N-O**) the HD alone. Transgenes were VCtagged derivatives. Rescue percentages show that HD rescues 80% of LT3 muscle alteration. In comparison, Ubx deleted from C- or N-terminal domains rescue 100% of the muscle alteration. n=8 embryos per genotype. Scale bar 50 µm. (**P-R**) Pull-down assay using (**P**) the indicated GST-fused Ubx derivatives and *Drosophila* S2R+ cell extracts expressing U1-70K-α. Input is indicated. (**Q-R**) Quantification of interactions relative to GST-$Ubx^{FL}$ full-length signal is indicated. n=3 biological replicates. Bars represent mean ±SEM. Statistics by one-way ANOVA (***P*< 0.01).

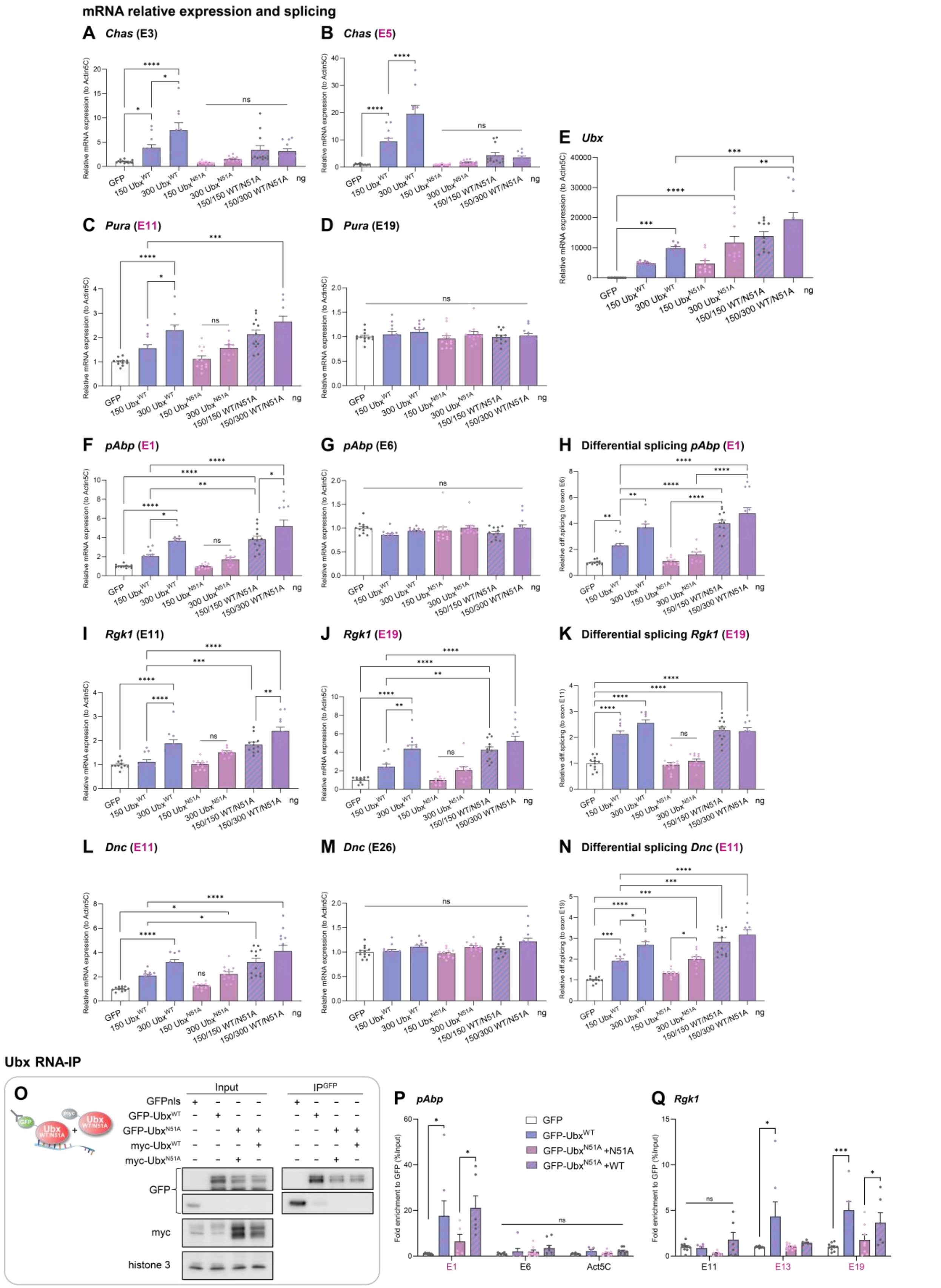


**Figure S2: Ubx$^{N51A}$ synergises with Ubx$^{WT}$ splicing activity and binds differentially spliced exons exclusively spliced.** (**A-N**) RTqPCR experiments showing the differential exon expression over *Actin5C* and (**H, K, N**) differential retention (Differential splicing) of exon cassettes (pink) over constitutive exons (black) for (**A-B**) *Chas,* (**C-D**) *Pura,* (**F-G**) *pAbp,* (**I-K**) *Rgk1,* (**L-N**) *Dnc,* in *Drosophila* S2R+ cells expressing GFP control (white), $Ubx^{WT}$ (blue), $Ubx^{N51A}$ (pink) or both co-expressed (purple). Transfected plasmid quantity is indicated (ng). (**E**) *Ubx* mRNA expression was controlled. This showed that $Ubx^{N51A}$ synergises with $Ubx^{WT}$ splicing activity on Ubx target genes only regulated at the splicing level (*pAbp*, *Dnc,* and *Rgk1* to a reduced extent). n=4 biological triplicates. (**O**) Western-blot control of Ubx-RNA immunoprecipitation. Input and IP-GFP are presented. (**P-Q**) RIP-RTqPCR performed in cells expressing GFP, GFP-$Ubx^{WT}$ and GFP-$Ubx^{N51A}$ with myc-$Ubx^{N51A}$ or myc-$Ubx^{WT}$ for constitutive and alternative exons of (**P**) *pAbp*, (**P**) *Actin5C* as control and (**Q**) *Rgk1*. Values are relative enrichment over GFP calculated as the percentage of input. (E+number)=exon, differentially spliced exons (pink). n=4 biological duplicates. Bars represent mean ±SEM. Statistics by one-way ANOVA (**P*< 0.05, ***P*< 0.01, ****P*< 0.001, *****P*< 0.0001, ns = non-significant).

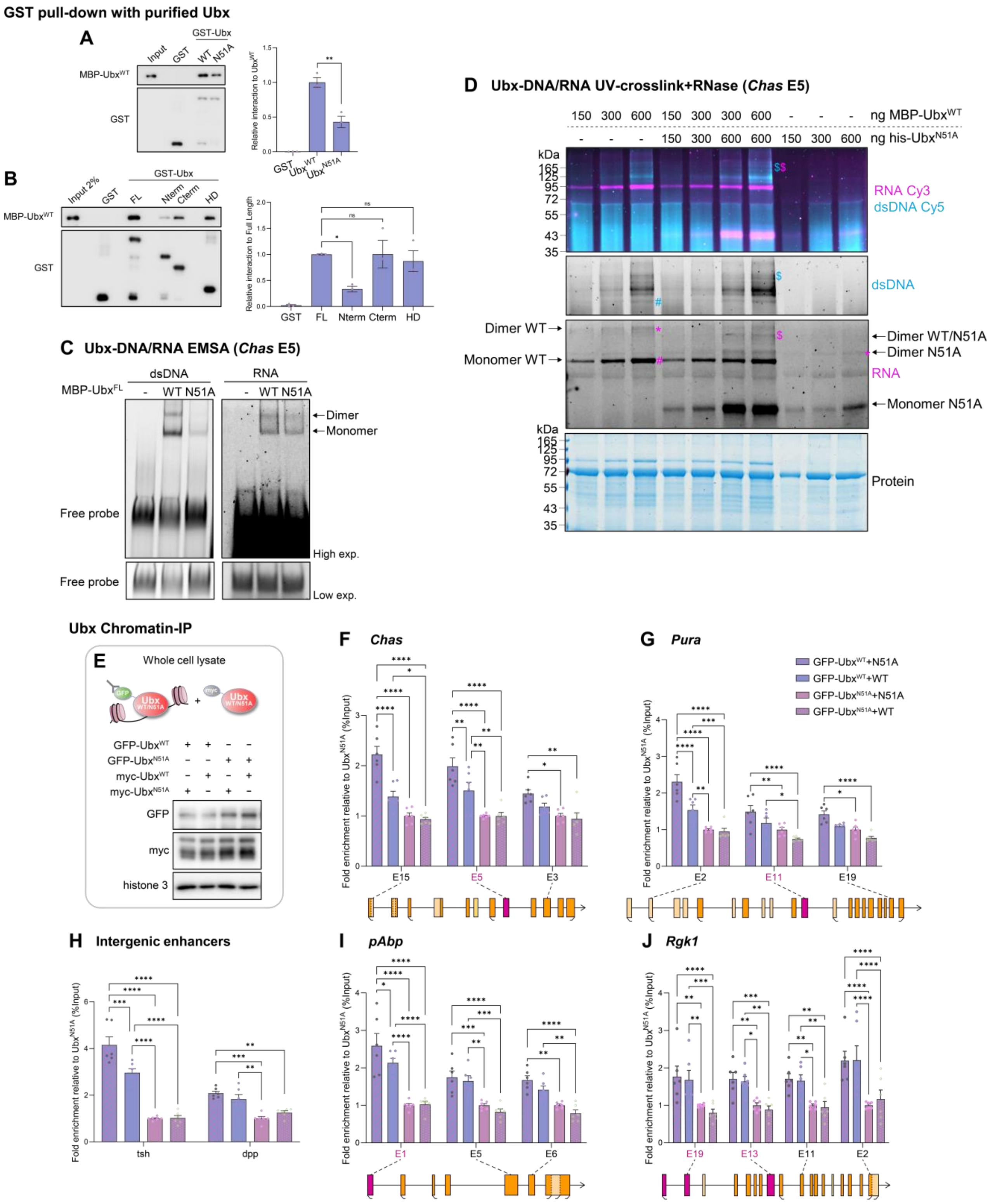


**Figure S3: Ubx homodimerizes on nucleic acids.** (**A-B**) Pull-down assay using the indicated GST-fused Ubx derivatives and purified his-MBP-Ubx proteins. Input is loaded as indicated. Quantifications of interactions relative to GST-Ubx$^{FL}$ full-length are indicated. n=3 biological replicates. Bars represent mean ±SEM. (**C**) Native electrophoretic mobility shift assay (EMSA) with purified proteins performed on dsDNA 3'Cy5-labelled and RNA 3'Cy3-labelled *Chas* exon E5 probes. Dimer and monomer are indicated. (**D**) Protein-RNA/dsDNA interaction followed by UV-crosslink and RNase digestion, performed *in vitro* with purified proteins his-MBP-Ubx$^{WT}$ and his-Ubx$^{N51A}$ as indicated. Interactions were detected on SDS-PAGE gels by Cy3-UTP signal for RNA (magenta), Cy5-labelled dsDNA (cyan) *Chas* exon E5 probes and stained by Coomassie to reveal the protein content (lower panel, BSA is detected at 70kDa). Molecular marker is indicated. Complexes with potential monomer Ubx$^{WT}$+DNA+RNA (#) or heterodimer Ubx$^{WT}$/Ubx$^{N51A}$+DNA+RNA ($) are indicated. Homodimers of Ubx$^{WT}$ or Ubx$^{N51A}$ on RNA are indicated (*). (**E**) Immunostaining control of expression from whole-cell lysate used for Chromatin-immunoprecipitation (ChIP). (**F-J**) ChIPqPCR experiments of GFP-Ubx$^{WT}$ or GFP-Ubx$^{N51A}$ co-expressed with myc-Ubx$^{WT}$ or myc-Ubx$^{N51A}$ are presented as a percentage of enrichment relative to input and to “N51A+N51A” condition (set to 1, pink). The Ubx binding on the proximal and distal exons to the Transcription Start Site (TSS) of (**F**) *Chas,* (**G**) *Pura,* (**I**) *pAbp,* (**J**) *Rgk1* is displayed relative to a schematic of the gene architecture as well as (**H**) intergenic Ubx-regulatory enhancers *dpp* and *tsh*. Differentially spliced exons are in pink. The transcription directionality is represented (black arrow). The alternative TSS and TTS are in brackets. n=3 biological duplicates. Notably, myc-Ubx$^{WT}$ competes with GFP-Ubx$^{WT}$ on the chromatin, reducing the Ubx-ChIP enrichment (blue). Bars represent mean ±SEM. Statistics test by one-way ANOVA (**P*< 0.05, ***P*< 0.01, ****P*< 0.001, *****P*< 0.0001, ns = non-significant).

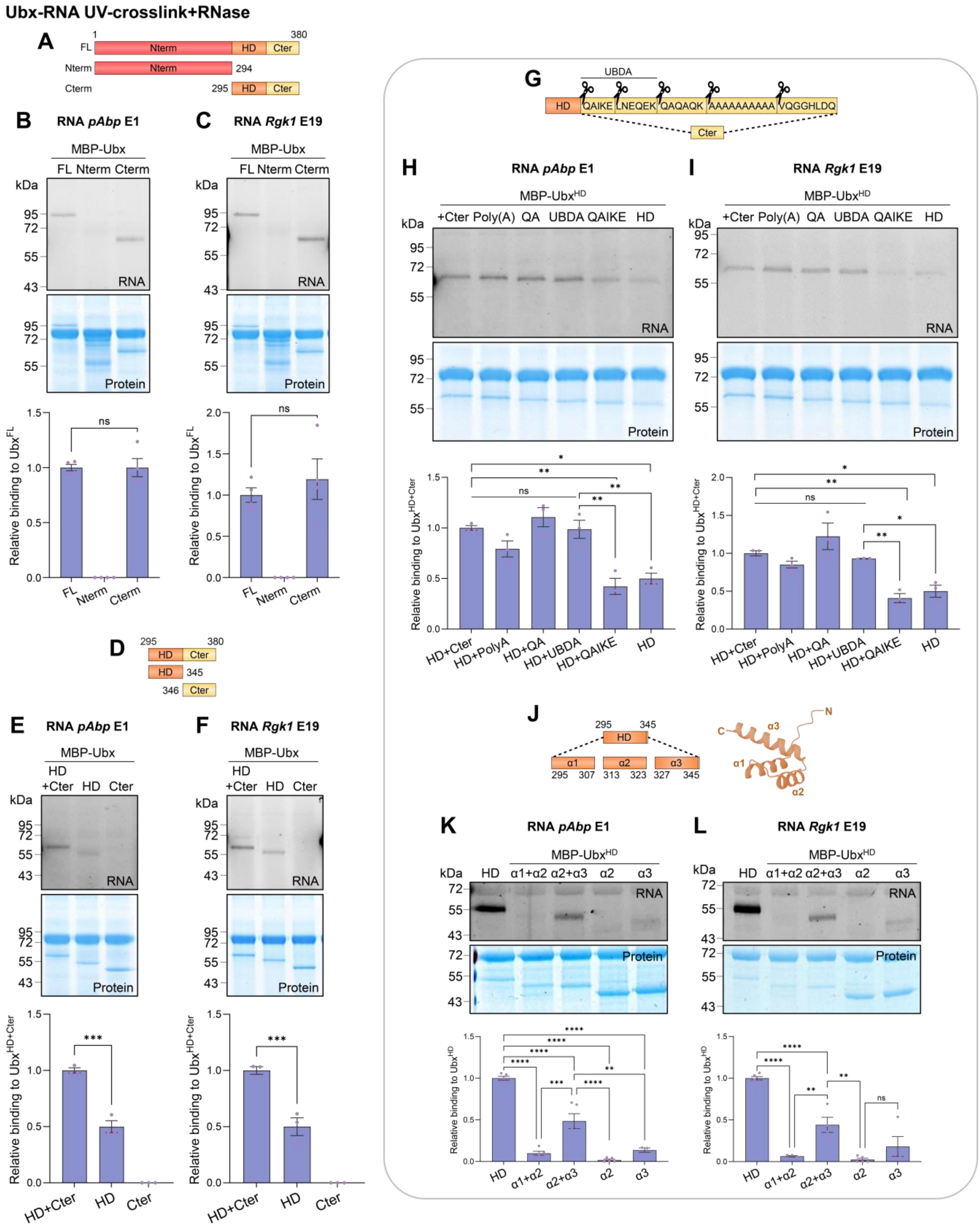


**Figure S4**: **The HD α3-helix is central for Ubx RNA binding, while the UBDA motif confers affinity.** (**A, D, G, J**) Cartoon of domains and motifs dissected by *in vitro* UV-crosslinking assays. (**B-C, E-F, H-I, K-L**) Protein-RNA interaction followed by UV-crosslink and RNase digestion performed in vitro with purified proteins his-MBP-Ubx derivatives as indicated on (**B, E, H, K**) *pAbp* exon E1 or (**C, F, I, L**) *Rgk1* exon E19 RNA probes. Cy3-UTP (RNA) signal detected interactions, Coomassie reveals the protein content (BSA is detected at 70kDa). Molecular marker is indicated. (**J**) HD structure generated with *AlphaFold*. Quantification of relative RNA-binding of Ubx derivatives compared to (**B-C**) full-length (FL), (**E-F, H-I**) HD+Cter or (**K-L**) HD for each RNA probe normalised to Coomassie staining. n=3 biological replicates. Bars represent mean ±SEM. Statistical test by one-way ANOVA (*P< 0.05, **P< 0.01, ***P< 0.001, ****P< 0.0001, ns = non-significant).

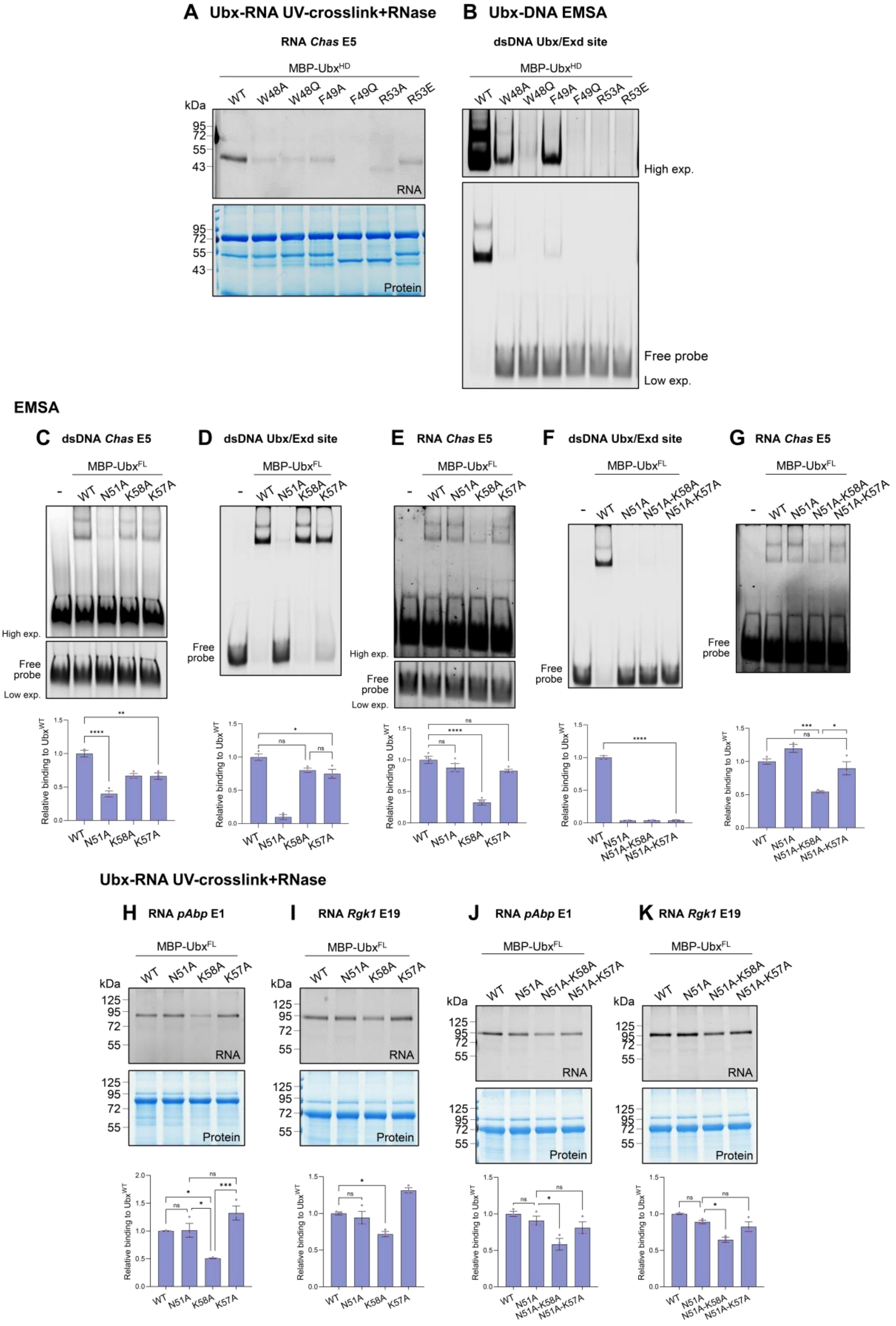


**Figure S5: The K58A mutation impairs Ubx-RNA binding but not DNA binding, while the K57A mutation does not alter Ubx-RNA binding *in vitro*.** (**A**) UV-crosslinking assays with purified proteins his-MBP-Ubx HD on *Chas* exon E5 RNA probes. Cy3-UTP signal (RNA) detected interactions on SDS-PAGE gels subsequently stained by Coomassie (Protein). BSA is detected at 70kDa. Molecular marker is indicated. Notably, F49Q and R53A mutations altered the structure of the *in vitro*-produced HD. (**B**) Native EMSA with purified proteins performed on 5'-Cy5-labelled dsDNA probes with optimal Ubx/Exd binding site. High and low exposures (exp.) of the supershift are presented. n=2 biological replicates. (**C-G**) Native EMSA with purified proteins performed from left to right on (**C**) 3'-Cy5-labelled dsDNA *Chas* E5, (**D, F**) 5'-Cy5-labelled dsDNA Ubx/Exd site, (**E, G**) 3'Cy3-labelled *Chas* E5 RNA probes for (**C-G**) $Ubx^{WT}$, $Ubx^{N51A}$ ($DNA^{mut}$), with (**C-E**) $Ubx^{K58A}$ ($RNA^{mut}$), $Ubx^{K57A}$ ($DNA^{+/-mut}$) and with (**F-G**) $Ubx^{N51A-K58A}$ ($DNA^{mut}$/$RNA^{mut}$) and $Ubx^{N51A-K57A}$ ($DNA^{mut}$). High and low exposures (exp.) of free probes are presented. (**H-K**) UV-crosslinking assays with purified proteins his-MBP-Ubx derivatives as indicated on (**H, J**) *pAbp* exon E1 and (**I, K**) *Rgk1* exon E19 RNA probes. This showed that the K58A mutation of the HD impacts Ubx-RNA binding ability *in vitro* but not the K57A mutation. Quantifications of relative RNA binding of Ubx derivatives compared to full-length (FL) proteins for distinct RNA probes normalised to Coomassie staining for UV-crosslinking assay and RNA/DNA binding by EMSA comparing the band shift to free probe and normalise to the WT protein. n=3 biological replicates. Bars represent mean ±SEM. Statistics by one-way ANOVA (*P< 0.05, **P< 0.01, ***P< 0.001, *****P*< 0.0001, ns = non-significant).

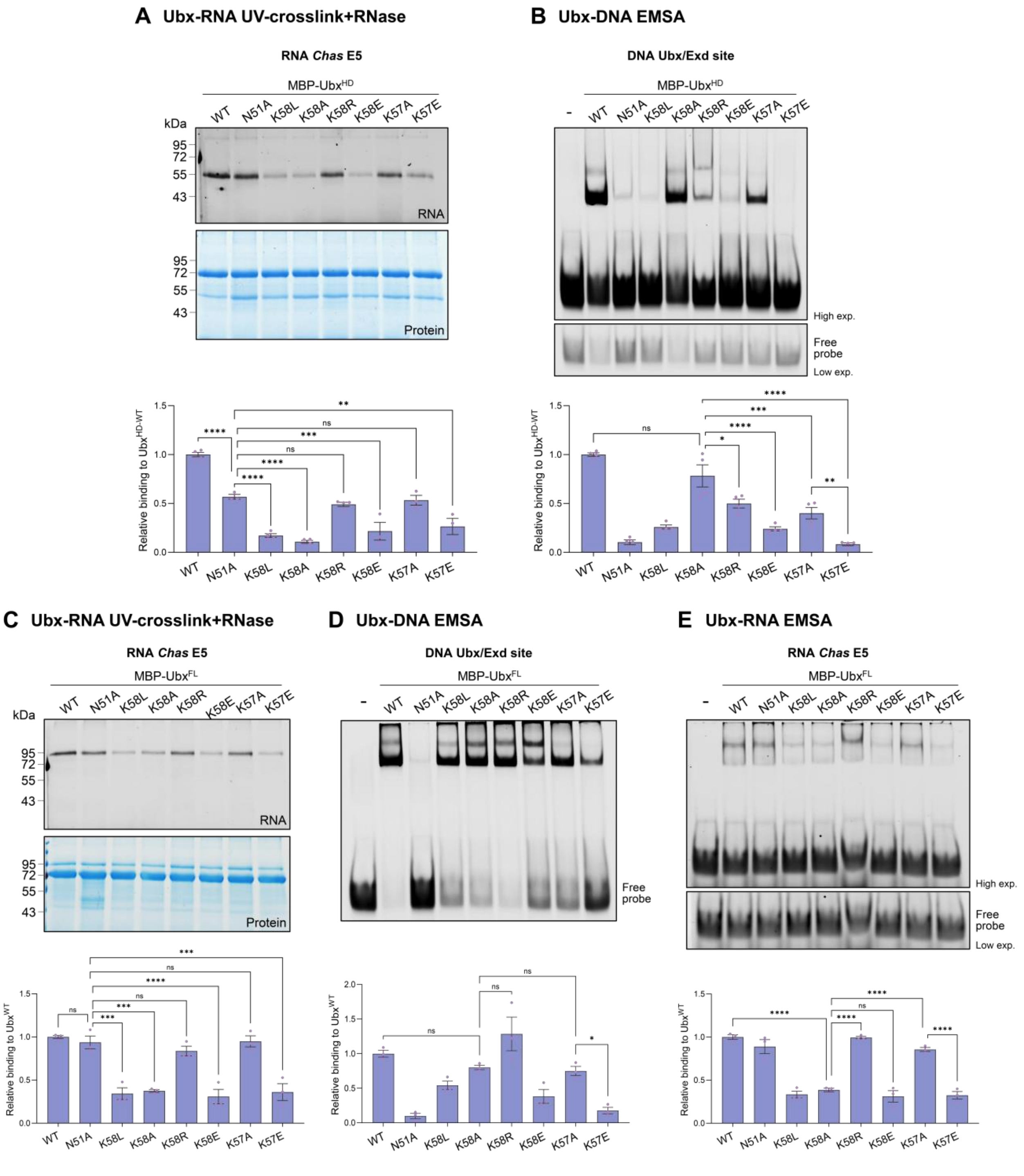


**Figure S6: Mutation of lysine K58 to alanine is optimal for impairing Ubx-RNA binding ability *in vitro*.** (**A**) UV-crosslinking assays with purified proteins his-MBP-Ubx derivative as indicated with HD (**A**) or full-length (FL) (**C**) protein and *Chas* exon E5 RNA probes. Interactions were detected on SDS-PAGE gels by Cy3-UTP signal (RNA) and stained by Coomassie (Protein). BSA is detected at 70kDa. Molecular marker is indicated. (**B**) Native EMSA with purified proteins performed on 5'-Cy5-labelled dsDNA probes containing an optimal Ubx/Exd binding site performed *in vitro* with purified proteins his-MBP-Ubx derivative as indicated whether for (**B**) HD or (**D**) full-length (FL). (**E**) Native EMSA with purified proteins performed on 3'-Cy3-labelled *Chas* exon E5 RNA probes performed *in*

*vitro* with purified proteins full-length (FL) his-MBP-Ubx derivatives. Quantifications of relative RNA-binding of Ubx derivatives compared to WT (HD or FL) MBP-fused proteins for *Chas* exon cassette E5 RNA probes and normalised to Coomassie staining for UV-crosslink assay (**A, C**) and RNA/DNA binding by EMSA comparing the band shift to free probe and normalise to the WT protein (**B, D-E**). For EMSA, low and high exposures (exp.) of free probes are presented. n=3 biological replicates. Bars represent mean ±SEM. Statistics by one-way ANOVA (*P< 0.05, **P< 0.01, ***P< 0.001, ****$P$< 0.0001, ns = non-significant).

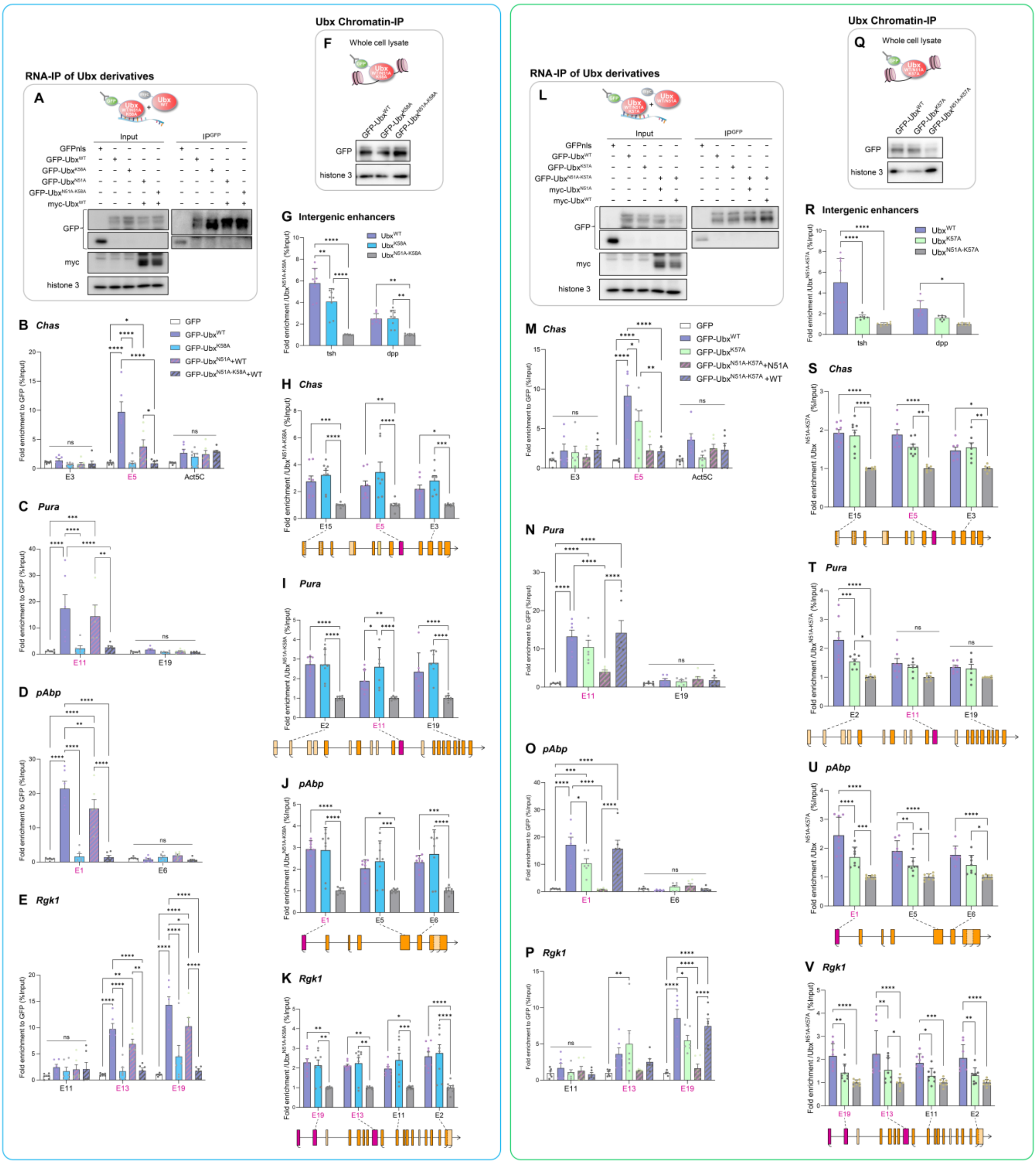


**Figure S7: The K58A mutation impairs Ubx-RNA binding *in vivo*.** (**A-E, L-P**) Ubx nuclear RNA-immunoprecipitation (**A, L**) western-blot control of input and IP-GFP are presented. (**B-E, M-P**) RIP-RTqPCR performed in cells expressing GFP, GFP-Ubx$^{WT}$, (**B-E**) GFP-Ubx$^{K58A}$, GFP-Ubx$^{N51A}$ with myc-Ubx$^{WT}$ and GFP-Ubx$^{N51A\text{-}K58A}$ with myc-Ubx$^{WT}$ and (**M-P**) GFP-Ubx$^{K57A}$, and GFP-Ubx$^{N51A\text{-}K57A}$ with myc-Ubx$^{WT}$ or myc-Ubx$^{N51A}$ on constitutive and alternative exons of (**B, M**) *Chas,* (**C, N**) *Pura*, (**D, O**) *pAbp* and (**E, P**) *Rgk1*. Values are relative enrichment over GFP calculated as the percentage of input. (E+number)=exon, differentially spliced exons (pink). (**B, M**) *Actin5C* mRNA is presented as a control.

n=3 biological duplicates. This showed that the K58A mutation impacts Ubx-RNA binding ability, mirroring the loss of splicing activity of $Ubx^{K58A}$ and $Ubx^{N51A\text{-}K58A}$ co-expressed with $Ubx^{WT}$. Similarly to $Ubx^{N51A}$, $Ubx^{N51A\text{-}K57A}$ (and $Ubx^{K57A}$) associates with differentially spliced exon RNA when co-expressed with $Ubx^{WT}$ *in vivo* in *Drosophila* cells (*Pura, pAbp, Rgk1*). This mirrors its synergistic capacity with $Ubx^{WT}$ splicing activity on exclusively differentially spliced genes. (**F, Q**) Immunostaining is a control of expression from whole-cell lysate used for Chromatin-immunoprecipitation (ChIP). (**G-K, R-V**) ChIPqPCR experiments of (**G-K**) GFP-$Ubx^{WT}$ (blue), GFP-$Ubx^{K58A}$ (light blue) or GFP-$Ubx^{N51A\text{-}K58A}$ (grey) are presented as a percentage of enrichment relative to input and to "N51A-K58A" condition (set to 1, grey). (**R-V**) ChIPqPCR experiments of GFP-$Ubx^{WT}$ (blue), GFP-$Ubx^{K57A}$ (green) or GFP-$Ubx^{N51A\text{-}K57A}$ (grey) are presented as a percentage of enrichment relative to input and to "N51A-K57A" condition (set to 1, grey). The Ubx binding on (**G, R**) intergenic Ubx-regulatory enhancers *dpp* and *tsh* and the proximal and distal exons to the Transcription Start Site (TSS) of (**H, S**) *Chas,* (**I, T**) *Pura,* (**J, U**) *pAbp,* (**K, V**) *Rgk1*. n=4 biological duplicates. The transcription directionality is represented. Differentially spliced exons are highlighted in pink. The alternative TSS and TTS are represented in brackets. This showed that $Ubx^{K58A}$ binds chromatin similarly to $Ubx^{WT}$ protein. Notably, $Ubx^{K57A}$ binding is strongly impaired on intergenic *tsh* enhancer and promoter-proximal exons of *Pura*, *pAbp* and *Rgk1*. Bars represent mean ±SEM. Statistical test by one-way ANOVA (*P< 0.05, **P< 0.01, ***P< 0.001, ****$P$< 0.0001, ns = non-significant).

**mRNA relative expression and splicing**

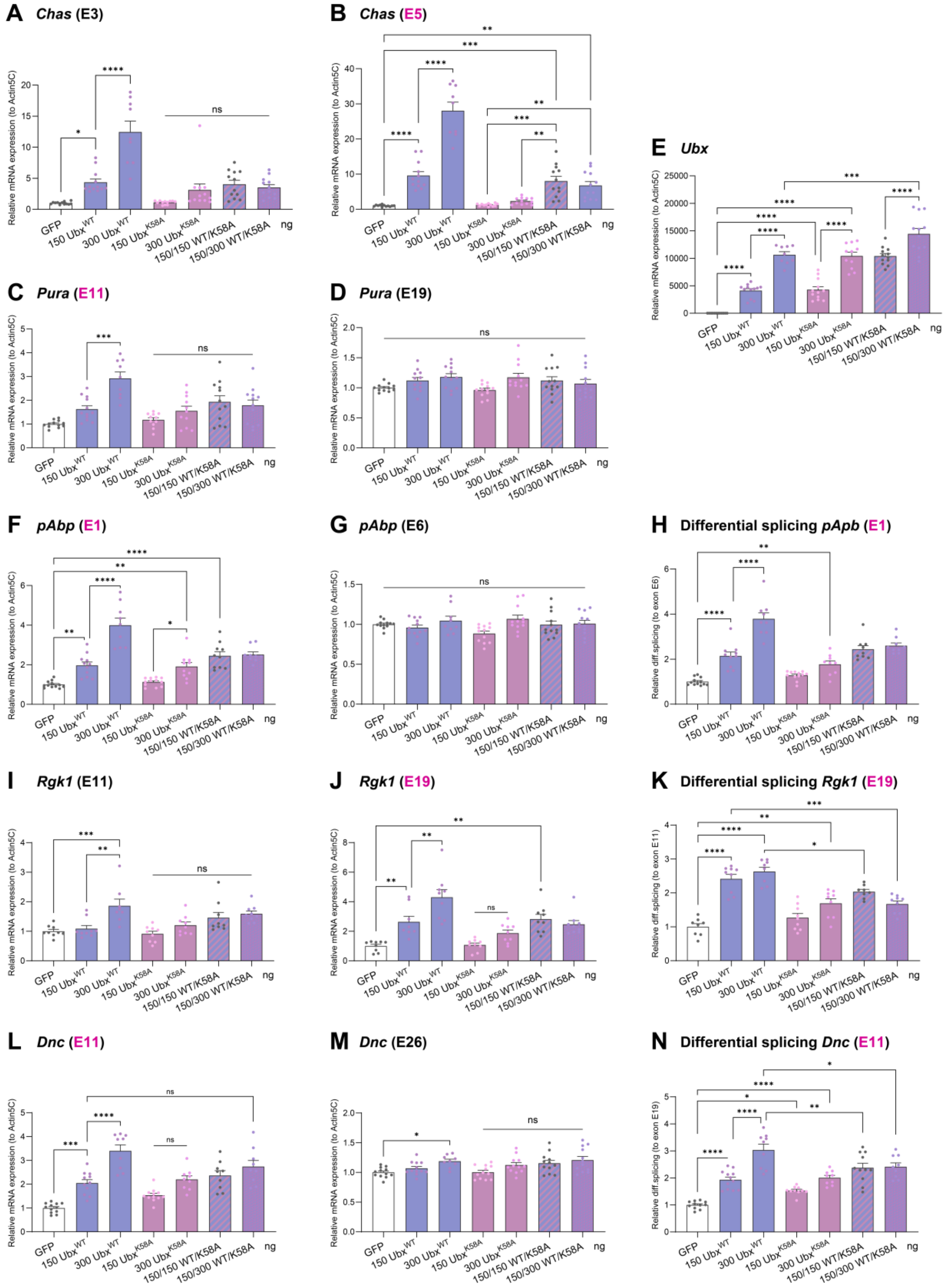


**Figure S8: The K58A mutation impairs Ubx splicing activity.** (**A-N**) RTqPCR experiments showing the (**A-B, C-D, F-G, I-J, L-M**) differential exon expression over *Actin5C* and (**H, K, N**) differential retention of exon cassettes (pink) over constitutive exons (Differential splicing) for (**A-B**) *Chas,* (**C-D**) *Pura,* (**F-H**) *pAbp,* (**I-K**) *Rgk1,* (**L-N**) *Dnc,* in *Drosophila* S2R+ cells expressing GFP control (white), $Ubx^{WT}$ (blue), $Ubx^{K58A}$ (pink) or both co-expressed (purple). Transfected plasmid quantity is indicated (ng). (**E**) *Ubx* mRNA expression was controlled. This showed that $Ubx^{K58A}$ ($RNA^{mut}$) failed to promote splicing or synergise with $Ubx^{WT}$ splicing activity on Ubx target genes. *n*=4 biological triplicates. Bars represent mean ±SEM. Statistical test by one-way ANOVA (*P< 0.05, **P< 0.01, ***P< 0.001, *****P*< 0.0001, ns = non-significant).

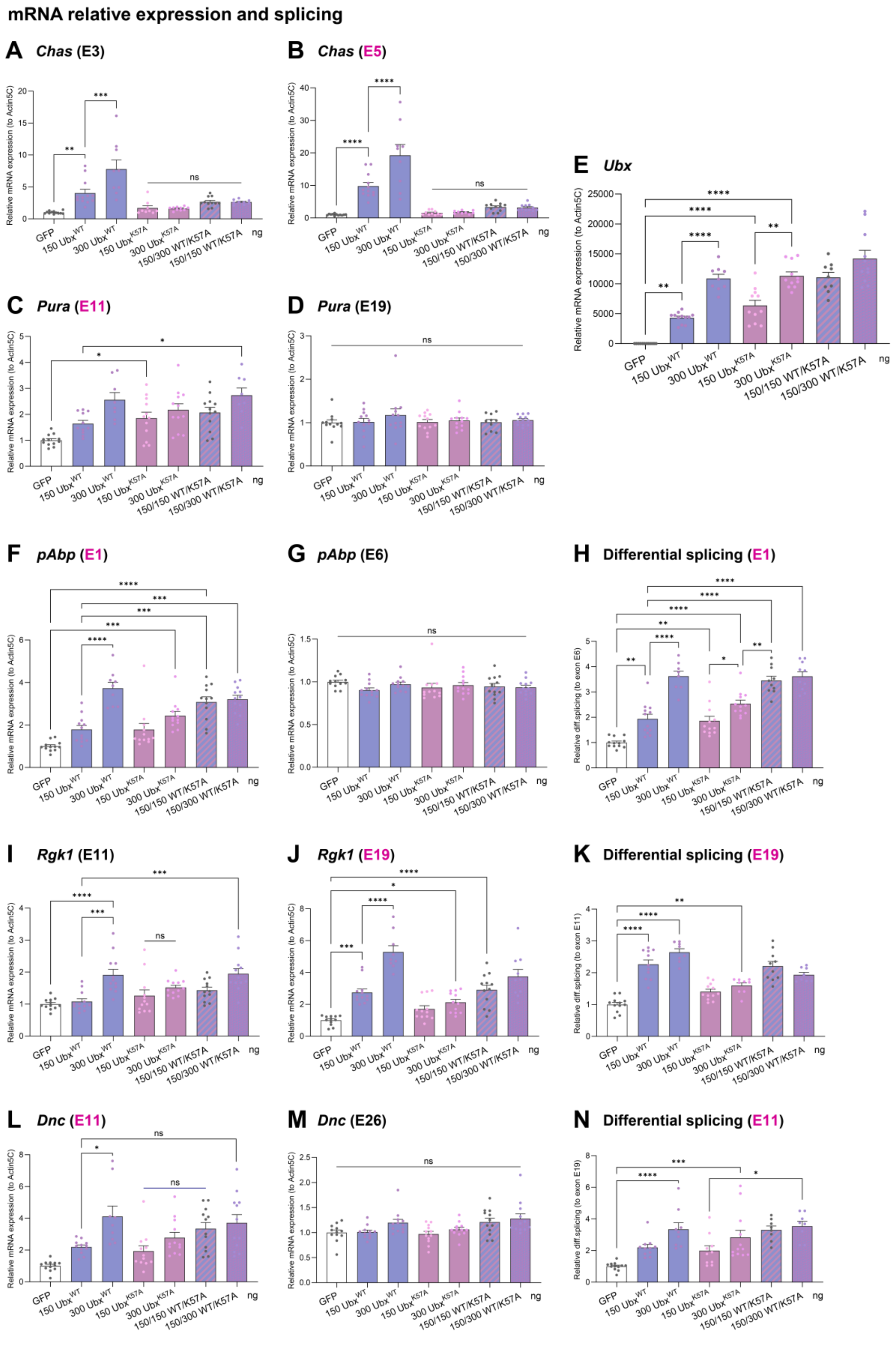


**Figure S9: Ubx$^{K57A}$ regulates splicing of Ubx-targets exclusively differentially spliced.** (**A-N**) RTqPCR experiments showing the (**A-B, C-D, F-G, I-J, L-M**) differential exon expression over *Actin5C*

and (**H, K, N**) differential retention of exon cassettes (pink) over constitutive exons (Differential splicing) for (**A-B**) *Chas,* (**C-D**) *Pura,* (**F-H**) *pAbp,* (**I-K**) *Rgk1,* (**L-N**) *Dnc,* in *Drosophila* S2R+ cells expressing GFP control (white), $Ubx^{WT}$ (blue), $Ubx^{K57A}$ (pink) or both co-expressed (purple). Transfected plasmid quantity is indicated (ng). (**E**) *Ubx* mRNA expression was controlled. This showed that $Ubx^{K57A}$ promotes differential splicing on Ubx target genes solely differentially spliced. n=4 biological triplicates. Bars represent mean ±SEM. Statistics by one-way ANOVA (*P< 0.05, **P< 0.01, ***P< 0.001, ****$P$< 0.0001, ns = non-significant).

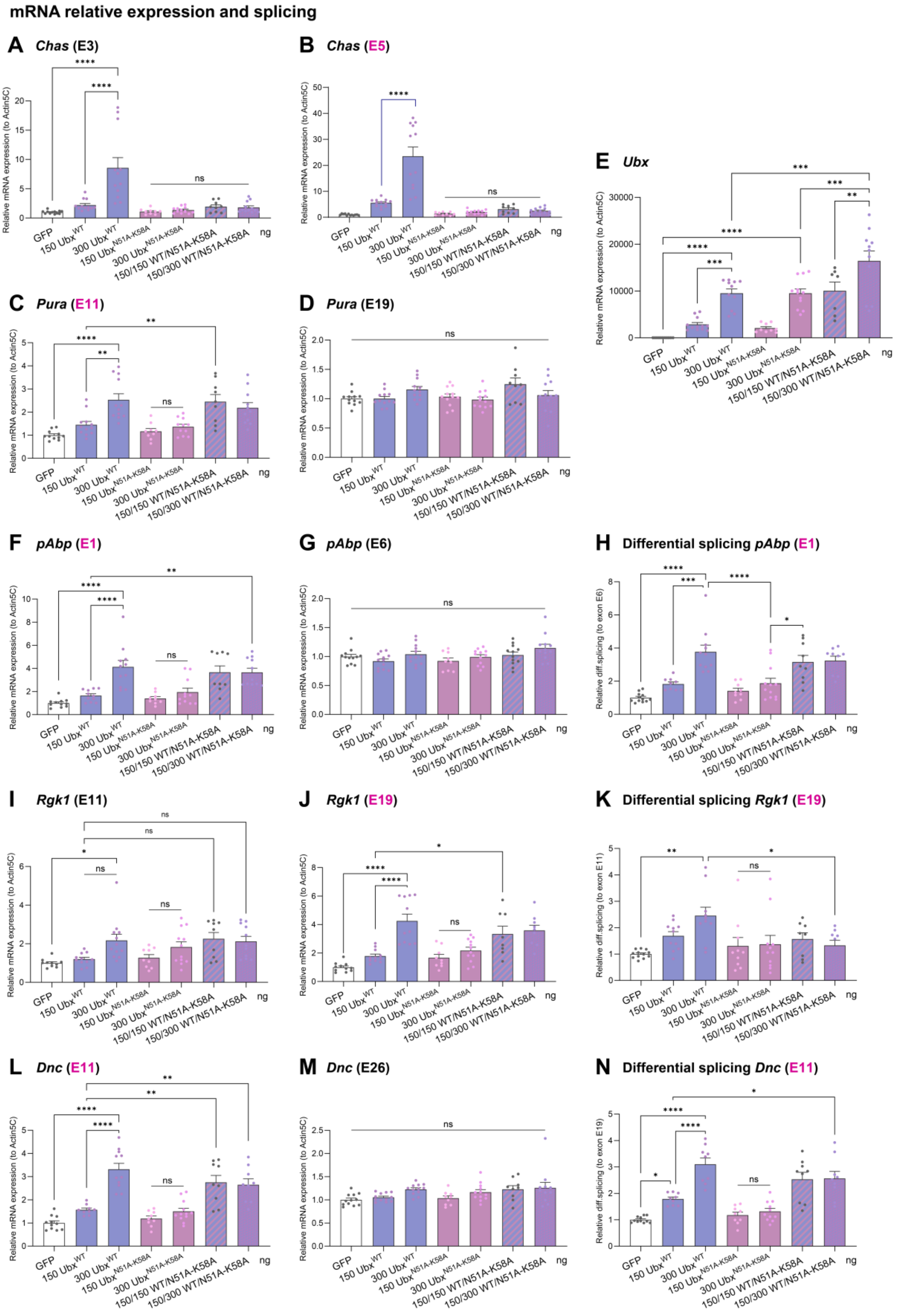


**Figure S10: The K58A mutation impairs Ubx$^{WT}$/Ubx$^{N51A}$ synergistic splicing activity.** (**A-N**) RTqPCR experiments showing the (**A-B, C-D, F-G, I-J, L-M**) differential exon expression over *Actin5C*

and (**H, K, N**) differential retention of exon cassettes (pink) over constitutive exons (Differential splicing) for (**A-B**) *Chas,* (**C-D**) *Pura,* (**F-H**) *pAbp,* (**I-K**) *Rgk1,* (**L-N**) *Dnc,* in *Drosophila* S2R+ cells expressing GFP control (white), $Ubx^{WT}$ (blue), $Ubx^{N51A\text{-}K58A}$ (pink) or both co-expressed (purple). Transfected plasmid quantity is indicated (ng). (**E**) *Ubx* mRNA expression was controlled. This showed that the K58A mutation circumvents the $Ubx^{WT}$/$Ubx^{N51A}$ synergistic splicing activity on Ubx target genes exclusively spliced. n=4 biological triplicates. Bars represent mean ±SEM. Statistics by one-way ANOVA (*P< 0.05, **P< 0.01, ***P< 0.001, ****$P$< 0.0001, ns = non-significant).

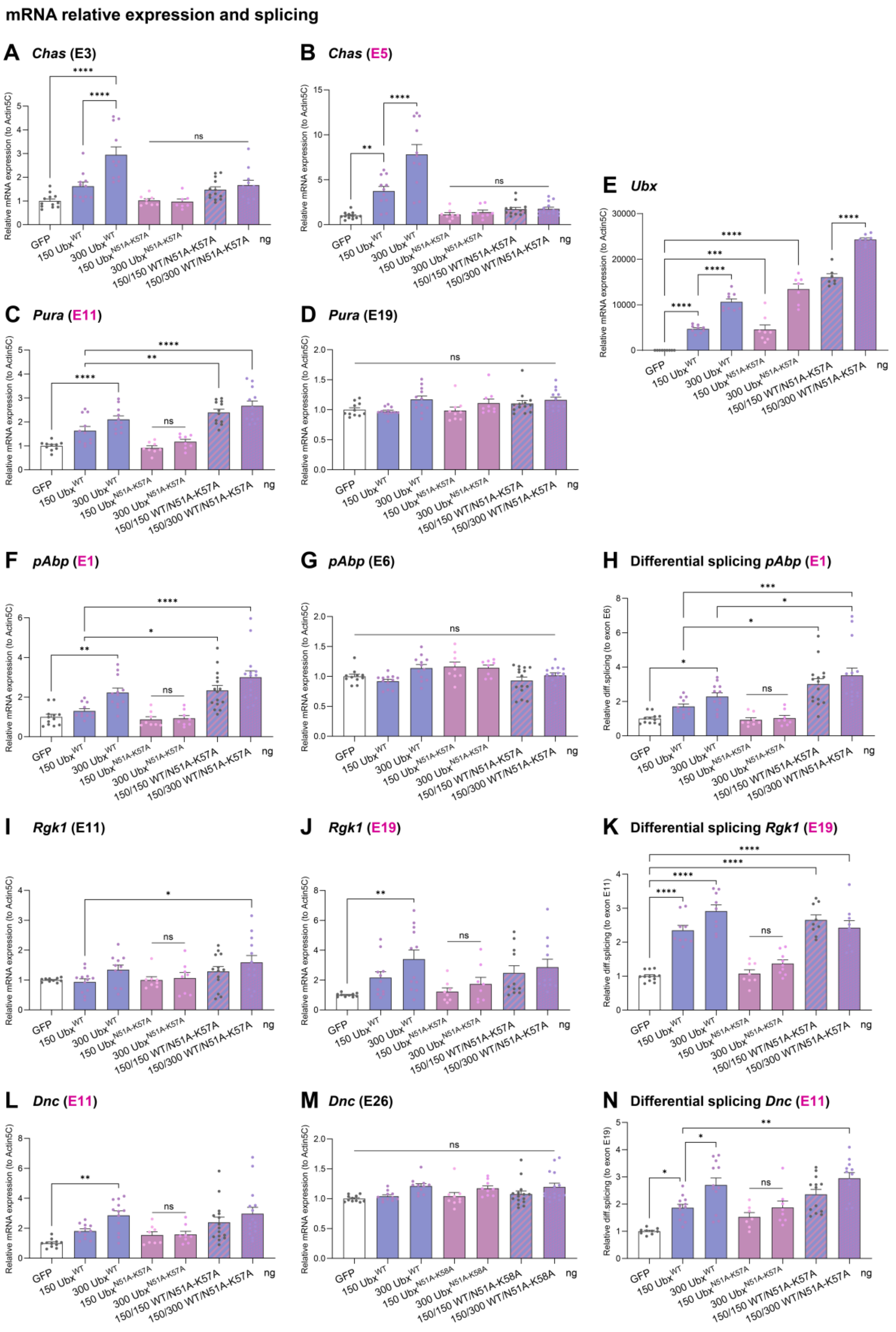


**Figure S11: The K57A mutation does not alter Ubx$^{WT}$/Ubx$^{N51A}$ synergistic splicing activity.** (**A-N**) RTqPCR experiments showing the (**A-B, C-D, F-G, I-J, L-M**) differential exon expression over

*Actin5C* and (**H, K, N**) differential retention of exon cassettes (pink) over constitutive exons (Differential splicing) for (**A-B**) *Chas,* (**C-D**) *Pura,* (**F-H**) *pAbp,* (**I-K**) *Rgk1,* (**L-N**) *Dnc,* in *Drosophila* S2R+ cells expressing GFP control (white), $Ubx^{WT}$ (blue), $Ubx^{N51A\text{-}K57A}$ (pink) or both co-expressed (purple). Transfected plasmid quantity is indicated (ng). (**E**) *Ubx* mRNA expression was controlled. This showed that the K57A mutation does not perturb the $Ubx^{WT}/Ubx^{N51A}$ synergistic splicing activity on Ubx target genes solely differentially spliced. n=4 biological triplicates. Bars represent mean ±SEM. Statistics by one-way ANOVA (*P< 0.05, **P< 0.01, ***P< 0.001, ****$P$< 0.0001, ns = non-significant).

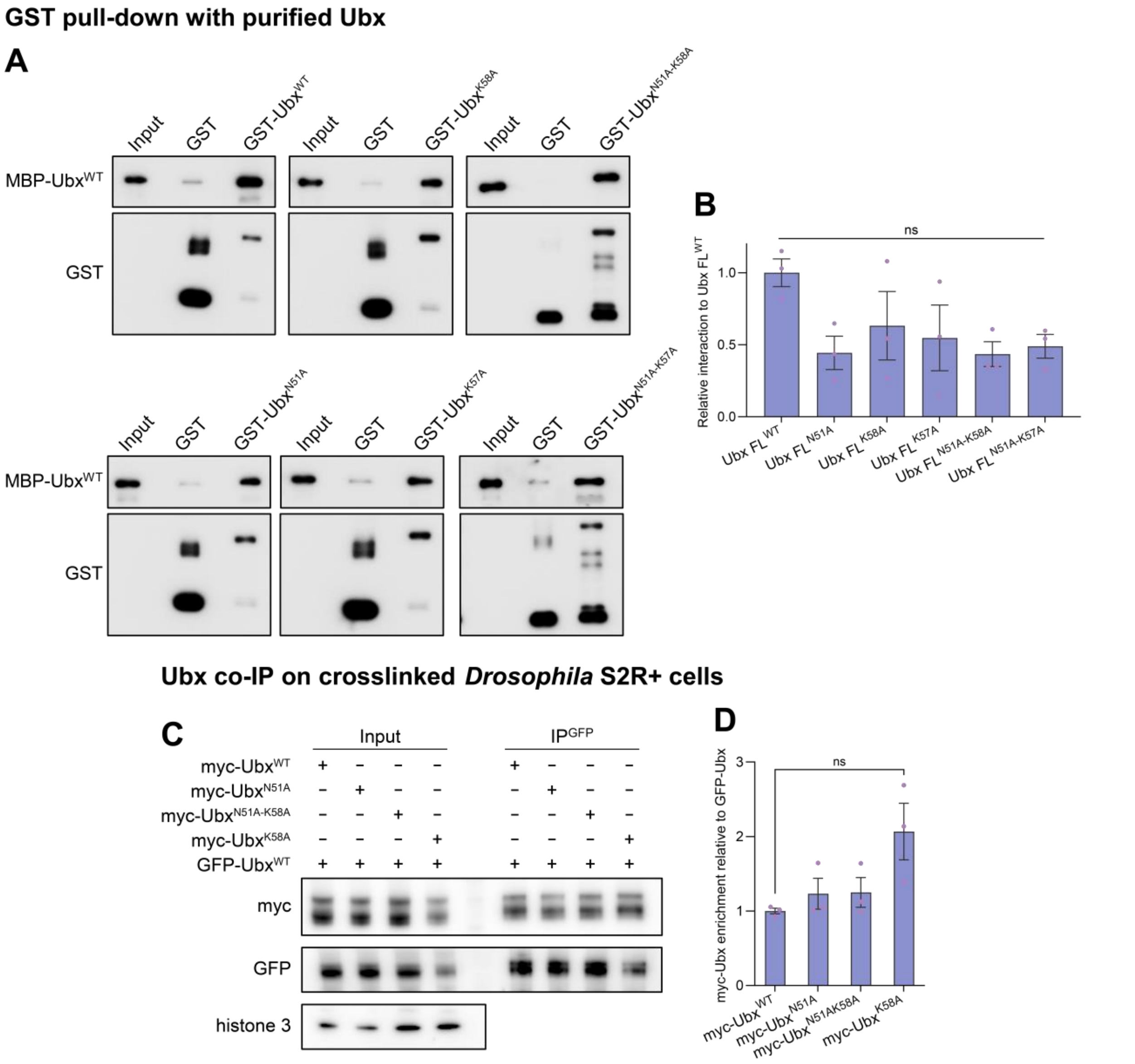


**Figure S12: The K58A does not alter Ubx dimerization potential *in vitro and in vivo*.** (**A-B**) Pull-down assay using the indicated GST-fused Ubx derivatives and purified his-MBP-Ubx proteins. Input is loaded as indicated. (**B**) Quantification of interactions relative to GST-Ubx$^{WT}$ and input signals is indicated. (**C-D**) Co-immunoprecipitation of myc-Ubx$^{WT}$, myc-Ubx$^{N51A}$, myc-Ubx$^{N51A\text{-}K58A}$ or myc-Ubx$^{K58A}$ with GFP-Ubx$^{WT}$ with crosslink. (**C**) Western blots were probed with the indicated antibodies. The input is shown as the control of expression and histone 3 as a loading control. (**D**) Quantification of relative enrichments of myc-Ubx relative to GFP-Ubx$^{WT}$ pull-down showed a robust interaction between Ubx$^{WT}$ and all mutants. n=3 biological replicates. Bars represent mean ±SEM. Statistical test by one-way ANOVA (ns = non-significant).

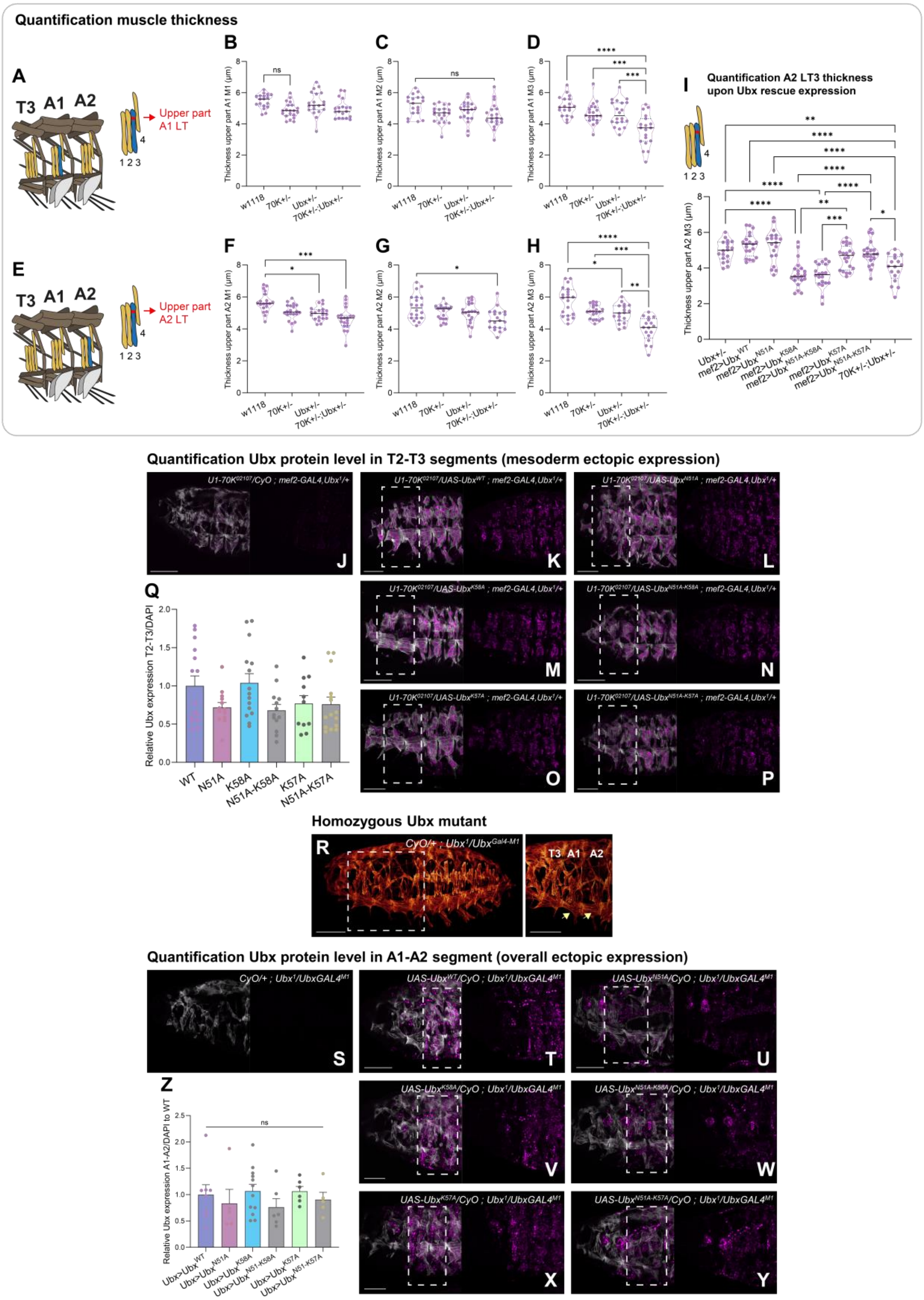


**Figure S13: Ubx-RNA binding activity contributes to muscle morphogenesis and homeotic functions.** (**A-I**) The rescue percentage was assessed by measuring the muscle thickness of the LT upper part of each (**B, F**) LT1, (**C, G**) LT2 and (**D, H**) LT3 muscle of abdominal segment (**A-D**) A1 and (**E-I**) A2. (**A, E**) Schematic of LT3 muscle (blue) and area quantified (red) are indicated. Notably, only the (**D, H**) LT3 showed a significant alteration of muscle thickness decrease in double heterozygous (*snRNP*)*U1-70K*$^{02107}$ with *Ubx*$^{1}$ embryos compared to *w*$^{1118}$, *U1-70K*$^{02107}$ or *Ubx*$^{1}$ single heterozygous mutant embryos. This has been observed in both (**B-D**) A1 and (**F-H**) A2 segments. (**I**) As observed for the A1 segment, pan-mesodermal expression of Ubx$^{WT}$, Ubx$^{N51A}$, Ubx$^{K57A}$ and Ubx$^{N51A\text{-}K57A}$ rescued the alteration of muscle thickness driven by the Ubx/U1-70K genetic interaction. All these constructs conserved their RNA binding ability. (**I**) As observed in the A1 segment, both Ubx-RNA binding mutant Ubx$^{K58A}$ and Ubx$^{N51A\text{-}K58A}$ were not able to rescue the thickness decrease of the A2 LT3 muscle. Muscle thickness was visualised with violin plots indicating median and quartiles. Statistics by one-way ANOVA (**P*< 0.05, ***P*< 0.01, ****P*< 0.001, *****P*< 0.0001, ns = non-significant). n=20 embryos. (**J-P**) is representative Z-slice of Ubx expression (magenta) in muscle (tropomyosin1, white) in rescue experiments of (**J**) *U1-70K*$^{02107}$ with *Ubx*$^{1}$ double heterozygous mutants with pan-mesodermal expression (*mef2-GAL4*) of (**K**) Ubx$^{WT}$, (**L**) Ubx$^{N51A}$, (**M**) Ubx$^{K58A}$, (**N**) Ubx$^{N51A\text{-}K58A}$, (**O**) Ubx$^{K57A}$ and (**P**) Ubx$^{N51A\text{-}K57A}$ transgenes. The area used for the Ubx quantification level is indicated with dashed lines and localised in T2-T3, where this is no endogenous Ubx expression. (**Q**) Graphical view represents Ubx intensity over DAPI for n=20 embryos. Notably, there is no significant difference in Ubx expression between the different Ubx transgenes. Scale bar 50 µm. (**R**) 3D projections of stage 16 embryos and zoom (tropomyosin1) of Ubx homozygous mutant *CyO,wg>LacZ/+;Ubx*$^{1}$*/Ubx*$^{Gal4\text{-}M1}$. Homeotic transformation of A1 and A2 into T3-like segments is highlighted (yellow arrow) and is the typical homozygous *Ubx* null mutant phenotype. (**S-Z**) Representative Z-slice of Ubx expression (magenta) in muscle (white) in rescue experiments of (**S**) Ubx homozygous mutants (*Ubx*$^{1}$*/Ubx*$^{Gal4\text{-}M1}$) with Ubx-like expression (*Ubx*$^{Gal4\text{-}M1}$) of (**T**) Ubx$^{WT}$, (**U**) Ubx$^{N51A}$, (**V**) Ubx$^{K58A}$, (**W**) Ubx$^{N51A\text{-}K58A}$, (**X**) Ubx$^{K57A}$ and (**Y**) Ubx$^{N51A\text{-}K57A}$ mutants. The area used for the Ubx quantification level is indicated with dashed lines and localised in A1-A2, where the Ubx identity-related function is pivotal. (**Z**) Graphical view represents Ubx intensity over DAPI for n=6-14 embryos. Notably, there is no significant difference in Ubx expression between the different Ubx transgenes. Scale bar 50 µm.

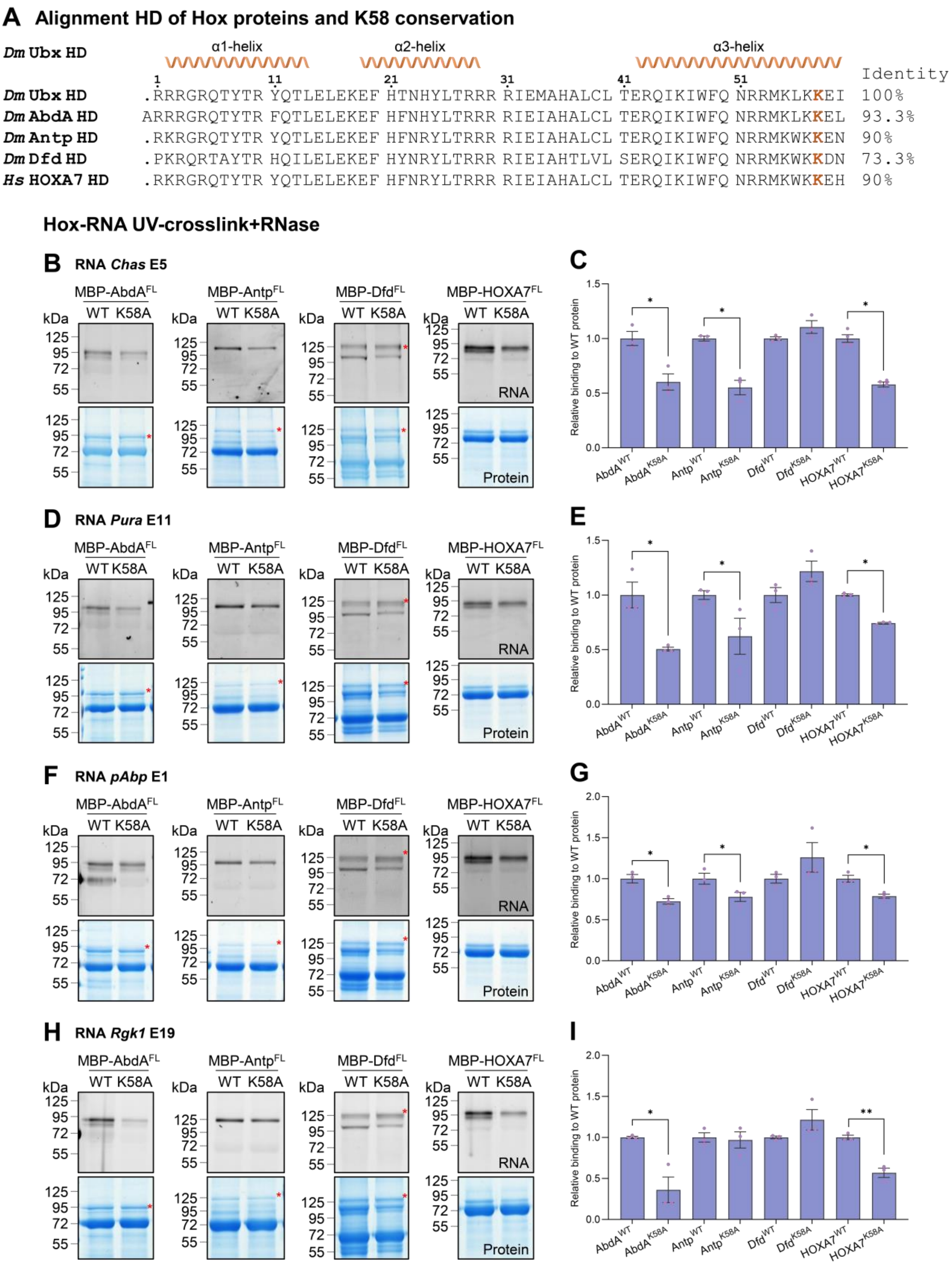


**Figure S14: RNA binding is a conserved feature of Hox proteins, with distinct potential binding interface K58-dependent or independent.** (**A**) HD domain alignment of *Drosophila* Hox proteins, Ubx, Deformed (Dfd, red star for specific band), Antennapedia (Antp), AbdominalA (AbdA) and the

Ubx-human ortholog HOXA7. The percentage of sequence matrix identity is provided. (**B-I**) Protein-RNA interaction by UV-crosslinking assay with purified proteins his-MBP-Ubx derivatives as indicated on (**B-C**) *Chas* exon E5, (**D-E**) *Pura* exon E11, (**F-G**) *pAbp* exon E1 and (**H-I**) *Rgk1* exon E19 RNA probes. Cy3-UTP signal (RNA) detected interactions on SDS-PAGE gels subsequently stained by Coomassie (Protein). BSA is detected at 70kDa. Molecular marker is indicated. (**C, E, G, I**) Quantification of relative RNA-binding of Hox derivatives compared to full-length WT his-MBP fused proteins for distinct RNA probes and normalised to Coomassie (Protein). This showed that RNA-binding is a conserved feature of the Hox proteins from *Drosophila* to Humans. Second, it revealed that while AbdA and human HOXA7 behave similarly to Ubx, the anterior Hox Antp and more strikingly Dfd, possess a distinct RNA binding interface as K58A mutation does not circumvent RNA-binding ability *in vitro*. n=3 biological replicates. Bars represent mean ±SEM. Statistical test by one-way ANOVA (**P*< 0.05, ***P*< 0.01).
